# Single-molecule detection of transient dimerization of opioid receptors 2: Heterodimer blockage reduces morphine tolerance

**DOI:** 10.1101/2024.07.25.605109

**Authors:** Peng Zhou, Rinshi S. Kasai, Wakako Fujita, Taka A. Tsunoyama, Hiroshi Ueda, Simone Pigolotti, Takahiro K. Fujiwara, Akihiro Kusumi

## Abstract

Heterodimerization of opioid receptors (ORs), MOR, KOR, and DOR, is implied in their functional regulation and diversification, and thus its understanding is crucial for developing better analgesic treatments. However, our knowledge on OR heterodimerization/heterodimers remains limited. Here, using single-molecule imaging and functional analysis, we found that MOR, the main morphine receptor, repeatedly forms *transient (≈250 ms) heterodimers* with DOR every 1-10 seconds, but not with KOR, whereas DOR and KOR also form transient heterodimers. We obtained all the heterodimer-monomer equilibrium constants and rate constants with/without agonists. We identified the critical heterodimer binding sites in the extracellular domains, in addition to the less-specific transmembrane domains, and developed soluble peptide blockers for MOR-DOR and DOR-KOR heterodimerization, using amino-acid sequences mimicking the extracellular binding sites. With these peptide blockers, we dissected the monomer/dimer roles in OR internalization and signaling. The soluble MOR-DOR heterodimer blocker reduced the development of long-term morphine tolerance in mice.

## Introduction

As elucidated in our companion paper,^1^ the transient homodimerization of μ-, κ-, and δ-opioid receptors (KOR, MOR, and DOR, respectively) provides a critical layer of regulation within the opioid signaling paradigm. New theories and methods for quantitative analyses of single-molecule imaging data developed there revealed that all ORs undergo transient homodimerizations for brief periods of 120-180 ms (*k*_off_ = 6.7-8.5 s^-1^), and they repeat dissociation and rebinding to another molecule every few seconds or less under physiological expression levels, with homodimer-dissociation equilibrium constants of 5.9-16.4 copies/µm^2^. The homodimerization is predominantly driven by unique 9-26 amino-acid stretches within the C-terminal cytoplasmic domain of each receptor, with less specific contributions from the transmembrane (TM) domains. Soluble but membrane-permeable peptides mimicking these C-terminal regions reduced homodimerization, enabling us to dissect the functions of homodimers. Compared to their monomeric counterparts, KOR-KOR and DOR-DOR homodimers (KK and DD homodimers)—unlike MOR-MOR homodimers (MM homodimers)—activate downstream signals differently in response to agonist binding, without affecting receptor internalization.

In this paper, we extend these advanced single-molecule imaging studies described in the companion paper to investigate OR *hetero*dimerization. The heterodimerization of three classical ORs introduces a new dimension to receptor interactions with G proteins, GRKs, and arrestins,^2, 3, 4, 5, 6^ providing additional opportunities for modulating OR functions. This modulation could be clinically utilized for managing chronic pain and treating neuropsychiatric disorders.^7, 8, 9, 10^

Given its crucial role in morphine response and the significant medical and social implications thereof, MOR has been the focus of extensive OR research,^10, 11, 12, 13, 14, 15^ including its heterodimerization with DOR and KOR (DM and KM heterodimers, respectively).^10, 13, 16, 17^ Extensive studies have reported DM heterodimerization^3, 18, 19, 20^ and downstream signals suggesting DM interactions,^21, 22, 23, 24^ although DM co-expression might be limited to small populations of neurons, such as excitatory interneurons, projection neurons in the spinal cord dorsal horn, and nociceptive neurons in dorsal root ganglia.^23, 25, 26^ In contrast, evidence for KM heterodimerization remains sparse, which suggests a dependence on the physiological context.^17, 27, 28^

The DK coupling exhibited unique signaling and functional regulation, making the DK heterodimer as another promising therapeutic target for pain treatment.^11, 29, 30^ Indeed, DK heterodimerization and observations suggesting DK interactions have also been reported,^11, 28, 29, 31, 32^ further highlighting its significance.

Despite these studies, the fundamental properties of heterodimers, including heterodimer-monomer dissociation equilibrium constants and rate constants (*K*_D_, *k*_off_, which is inverse lifetime, and *k*_on_), heterodimer interaction sites, and cellular- and tissue-level functions, remain enigmatic. This research addresses these fundamental issues by leveraging the advanced single-molecule image analysis described in the companion paper. Such understanding would form the basis for developing opioid analgesics with heightened efficacy and minimized tolerance.

The heterodimer-monomer equilibrium constant, *K*_D_ (hetero-*K*_D_), can only be obtained after the homodimer-monomer equilibrium constants *K*_D_s (homo-*K*_D_s) for the two constituent ORs are evaluated, as detailed in the companion paper. We also compare the properties and dimerization sites between homo- and hetero-dimers of ORs.

This study lays the groundwork for future explorations of dimerization phenomena across the broader GPCR family.^33, 34^ Heterodimerization among class-A GPCRs has been increasingly observed, offering insights into novel signaling pathways and pharmacological profiles.^7, 8, 16, 35, 36^ Examples include heterodimers between MOR and V1b vasopressin receptor,^37^ MOR and somatostatin receptor 2,^38^ dopamine D2 receptor and neurotensin NTS1 receptor,^39^ angiotensin II AT1 receptor and norepinephrine α2C-adrenergic receptor,^35^ cannabinoid receptors CB1 and CB2,^40^ dopamine receptors D1 and D2,^41^ and others, marking an emergent frontier in GPCR research with profound implications for drug discovery.

Here, we unequivocally demonstrate that, under low physiological expression conditions, DOR-MOR and DOR-KOR form metastable heterodimers, whereas MOR-KOR heterodimers are not detectable. These findings have been substantiated through examinations conducted with a level of quantification unparalleled in the existing biomedical literature, with the exception of the OR homodimer research described in our companion paper.

Furthermore, we discovered that the extracellular N-terminal domain interactions and extracellular loop 3 (EL3) interactions are specifically responsible for DM and DK heterodimerizations, respectively. In contrast, the TM domain interactions contribute less specifically, challenging the common belief that GPCR dimerizations are primarily mediated by TM domain interactions.^3, 16, 30, 42, 43, 44^ In addition, allosteric conformational changes involving both extracellular domains and TM domains might be involved.

Building on these findings, we demonstrated that soluble peptides mimicking the extracellular amino-acid sequences implicated in DM and DK heterodimerizations effectively inhibit heterodimer formation. In murine models, the administration of a soluble peptide blocker targeting the DM heterodimers into the cerebral ventricles diminishes the development of long-term morphine tolerance. These results strongly suggest that DM heterodimerization based on the N-terminal domain sequence occurs in neuronal tissues. These findings underscore potential avenues for improving opioid drug administration strategies aimed at mitigating tolerance and dependence.

## Results

### Metastable DM and DK, but not MK, heterodimers are detected in the PM

SNAPf-ORs and OR-Halos were expressed in CHO-K1 cells, which do not express ORs (Supplementary Fig. 1a).^45^ These tagged ORs are functional (Supplementary Fig. 1a, b) and labeled with SNAP-Surface 549 and Halo7-SaraFluor650T dyes. Their fluorescence characteristics are summarized in Supplementary Fig. 1c-e. The labeled OR molecules were simultaneously observed in two colors at the single molecule level at video rate (30 Hz) at 37°C, using a home-built total internal reflection fluorescence (TIRF) microscope. The cells expressing each OR species at fluorescence spot densities between 0.5-1 spot/µm^2^ (total number densities between 1-2 spots/µm^2^) after fluorescence labeling were selected for microscope observations. Virtually all of the OR fluorescent spots exhibited diffusion in the PM.

When DOR and MOR were expressed in the same cell, they exhibited frequent brief colocalization and co-diffusion, suggesting metastable DM heterodimer formation (Supplementary Videos 1 and 2; Fig. 1a). For quantifying colocalization, we employed a pair cross-correlation function (PCCF; Supplementary Fig. 1f; also see companion paper), which provides both a simple measure for heterodimer formation, the colocalization index, and a fundamental constant, the heterodimer-monomer dissociation equilibrium constant *K*_D_ (see Supplementary Note 2, Supplementary Fig. 2e, and related main text of the companion paper). As a control for incidental fluorescent spot colocalizations, the video frames in the green channel were rotated 180° before overlaying (see Supplementary Fig. 2d of the companion paper).

**Fig. 1:**
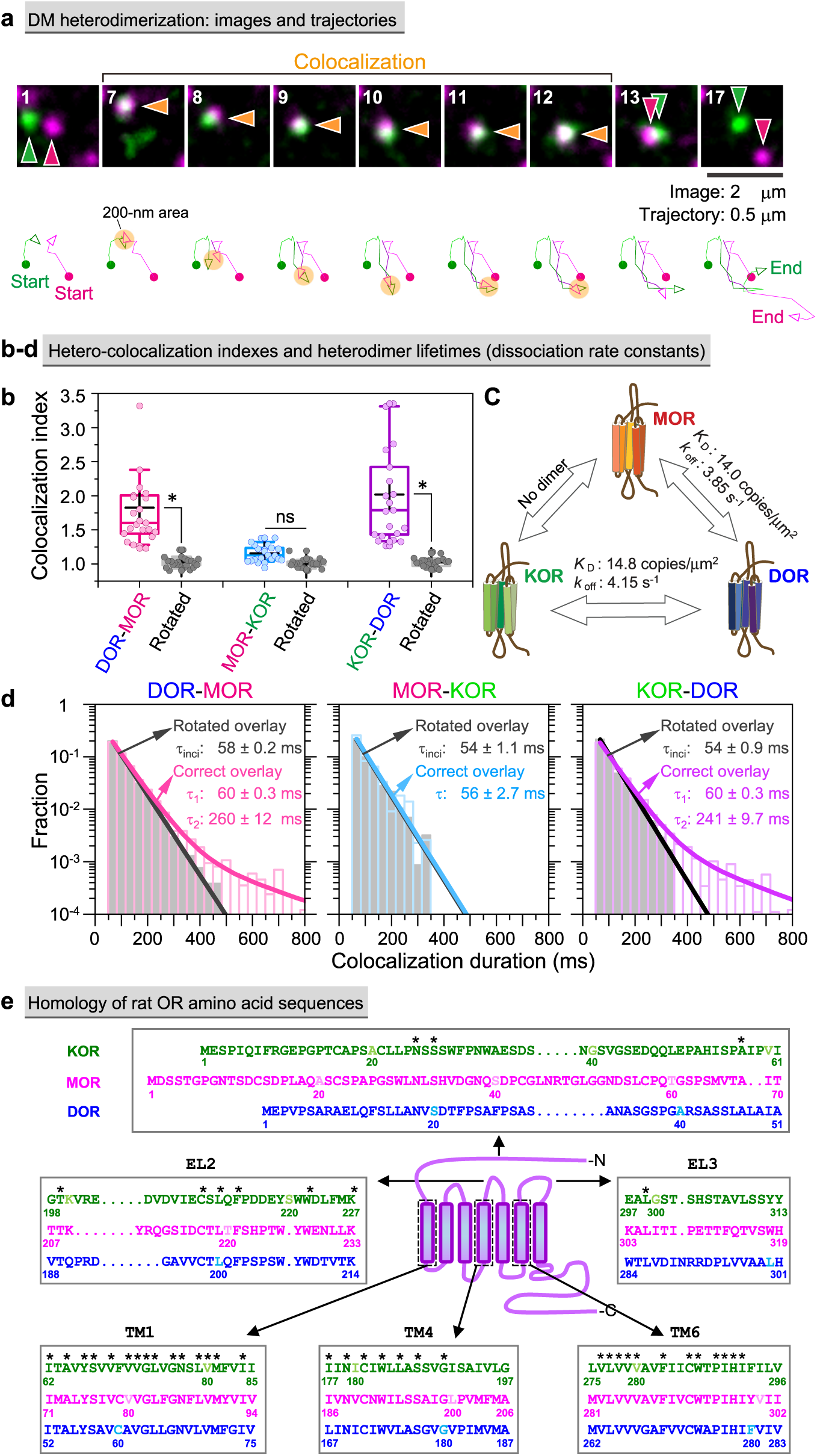
Transient DM and DK heterodimers, but not MK heterodimers, form in the PM. **a** Typical image sequence of simultaneous two-color single fluorescent molecule observations and trajectories of the molecules shown in the images. These molecules undergo transient hetero-colocalization and co-diffusion (SNAP-Surface 549-labeled SNAPf-DOR, green; SaraFluor650T-labeled MOR-Halo, magenta). **b** DM and DK heterodimers form, but MK heterodimers do not, as detected by colocalization indexes for correct and rotated overlays (see Supplementary Fig. 2c-e in the companion paper.^1^ Throughout this report, we employ the following conventions. For box plots, horizontal bars, crosses, boxes, and whiskers indicate the median values, mean values, interquartile ranges (25-75%), and 10-90% ranges, respectively; for bar graphs and plots showing mean values, SEM values are provided by bars; and in statistical analyses, * and ns (not significant) indicate *p* < 0.05 and *p* ≥ 0.05, respectively, obtained by Tukey’s multiple comparison test. The result set used for multiple comparison is indicated by the group of lines in each figure, and different sets are indicated by different colors (e.g., Fig. 2c, 3c and 7e). All statistical parameters (number of replicates and *p* values) are summarized in Supplementary Table 1. **c** Summary of *K*_D_ and *k*_off_ values for OR heterodimers. **d** Histograms showing the distributions of hetero-colocalization *durations* for correct and rotated overlays. The control histograms for rotated overlays (grey) were fitted by single exponential functions (black), providing the lifetimes of the incidental overlap events between the magenta and green spots (1_inci_). The histograms for correct overlays (colors) were fitted by the sum of two exponential functions: The faster decay time (τ_1_) was close to the lifetime of incidental overlaps (1_inci_), and the slower decay time provided the heterodimer lifetime (τ_2_). The heterodimer lifetime after correction for the trackable duration lifetime is shown in each box. The *k*_off_ values (1/τ_2_) calculated from τ_2_ are shown in **c**. **e** Comparison of the amino acid sequences among the three classical ORs (rat) in the domains where the amino acid homologies are lower. Asterisks mark the identical amino acids.

The colocalization index clearly indicated that DOR and MOR form DM heterodimers (Fig. 1b) and DOR and KOR form DK heterodimers (Fig. 1b and Supplementary Fig. 1f) in the PM (all statistical parameters including *p* values and numbers of replicates are summarized in Supplementary Table 1). However, at the expression levels employed in this study, KM heterodimers were not detected (Fig. 1b and Supplementary Fig. 1f), consistent with the previous result.^28^ The heterodimer-monomer dissociation equilibrium constants *K*_D_’s were obtained from the PCCFs (Fig. 1c).

The heterodimer lifetime (*1*_2_) was determined from *k*_off_ (summarized in Fig. 1c), evaluated from the distribution of colocalization *durations* (Fig. 1d; see Note S1 in the companion paper). The *1*_2_ values are 260 ± 12 ms for DM heterodimers and 241 ± 10 ms for DK heterodimers, after corrections for trackable duration lifetimes (Fig. 1d and Supplementary Fig. 1e). The histogram for the MK colocalization durations did not exhibit any *1*_2_ component (the decay component representing the heterodimer loss), consistent with the lack of detectable heterodimers for this pair (only incidental colocalization with a lifetime of *1*_1_; Fig. 1d).

To identify the amino-acid sequences responsible for specific heterodimerization, we initially examined the OR domains with lower amino-acid identities/homologies among the three ORs. The candidate sites we initially selected are summarized in Fig. 1e. The amino-acid sequence homologies in the cytoplasmic C-terminal domains are lower, but they are not included in this figure as they are involved in KK, MM, and DD homodimerizations (companion paper) and we unequivocally showed that they are not involved in DK and DM heterodimerizations (Fig. 2a and 3a).

**Fig. 2:**
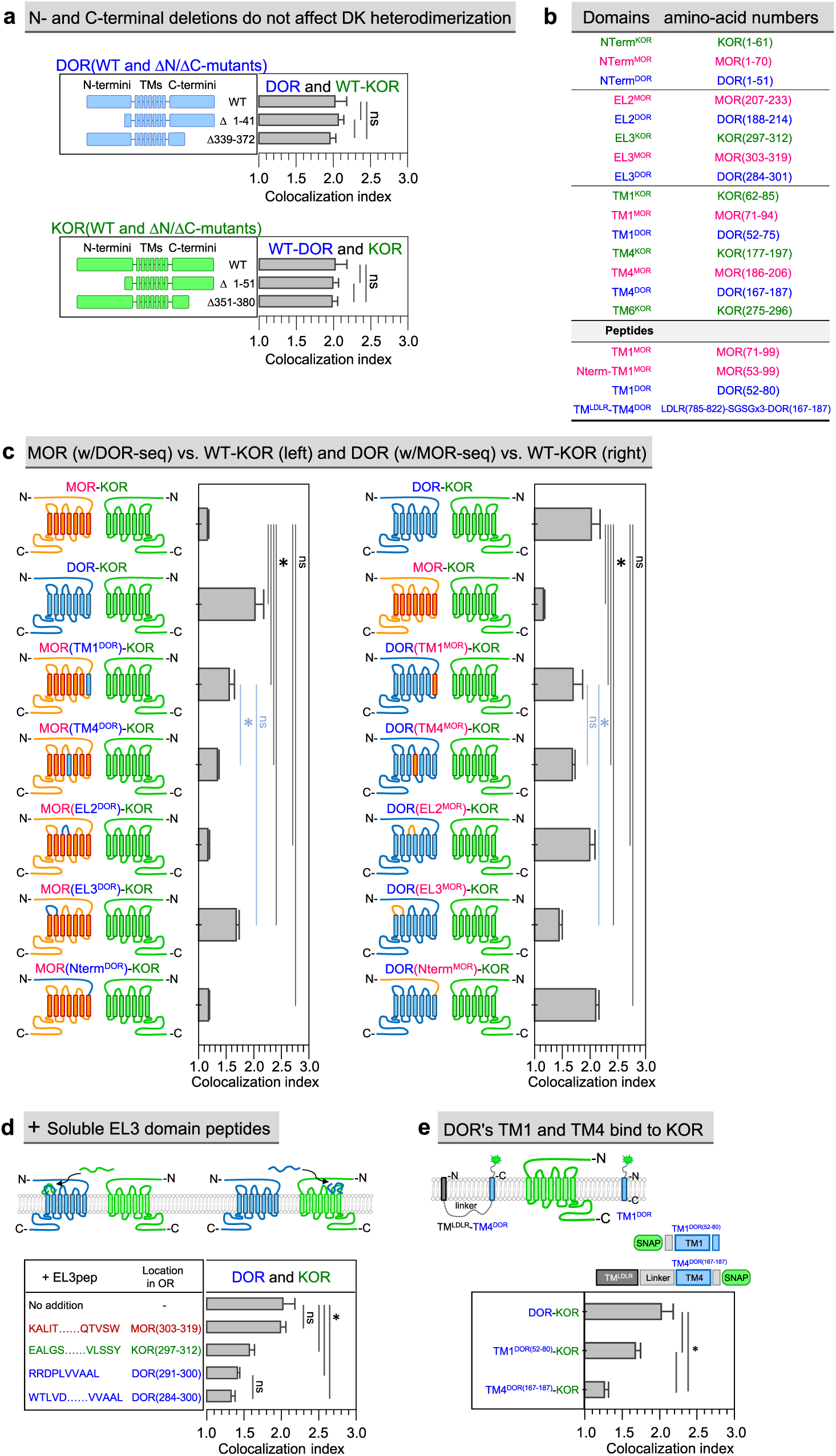
Interaction between EL3 domains of DOR and KOR is involved in DK heterodimerization, and DOR’s TM1 and TM4 are also involved. **a** The N- and C-terminal domains of DOR and KOR are not involved in DK heterodimerization. DK colocalization indexes of N/C-terminal deletion mutants of DOR and full-length KOR (top) and N/C-terminal deletion mutants of KOR and full-length DOR (bottom). **b** Summary of the names and exact amino-acid ranges of the OR domains and TM1-based peptides employed in this work. **c** DOR’s EL3, TM1, and TM4, but not EL2 and N-term, are involved in DK heterodimerization. (Left panel) Because MK heterodimerization scarcely occurs (top), we examined whether MK heterodimerization (MK colocalization index) is enhanced by substituting various MOR domains with the corresponding DOR domains, using WT-KOR as the binding partner (DK heterodimerization as a positive control; second). (Right panel) Because DOR binds to KOR to form DK heterodimers (top) and MK heterodimerization rarely occurs (second), we examined whether DK heterodimerization (DK colocalization index) is reduced by substituting various DOR domains with the corresponding MOR domains, using WT-KOR as the binding partner. **d** Both EL3 domains of DOR and KOR are involved in DK heterodimerization. Soluble peptides (1 µM) with the aa sequences of the EL3 domains (EL3-domain peptides) of DOR and KOR, but not that of MOR, block DK heterodimerization. **e** DOR’s TM1 and TM4 bind to KOR. Colocalization indexes show that the affinities to WT-KOR are greater in the order of WT-DOR > DOR’s TM1 > DOR’s TM4.

**Fig. 3:**
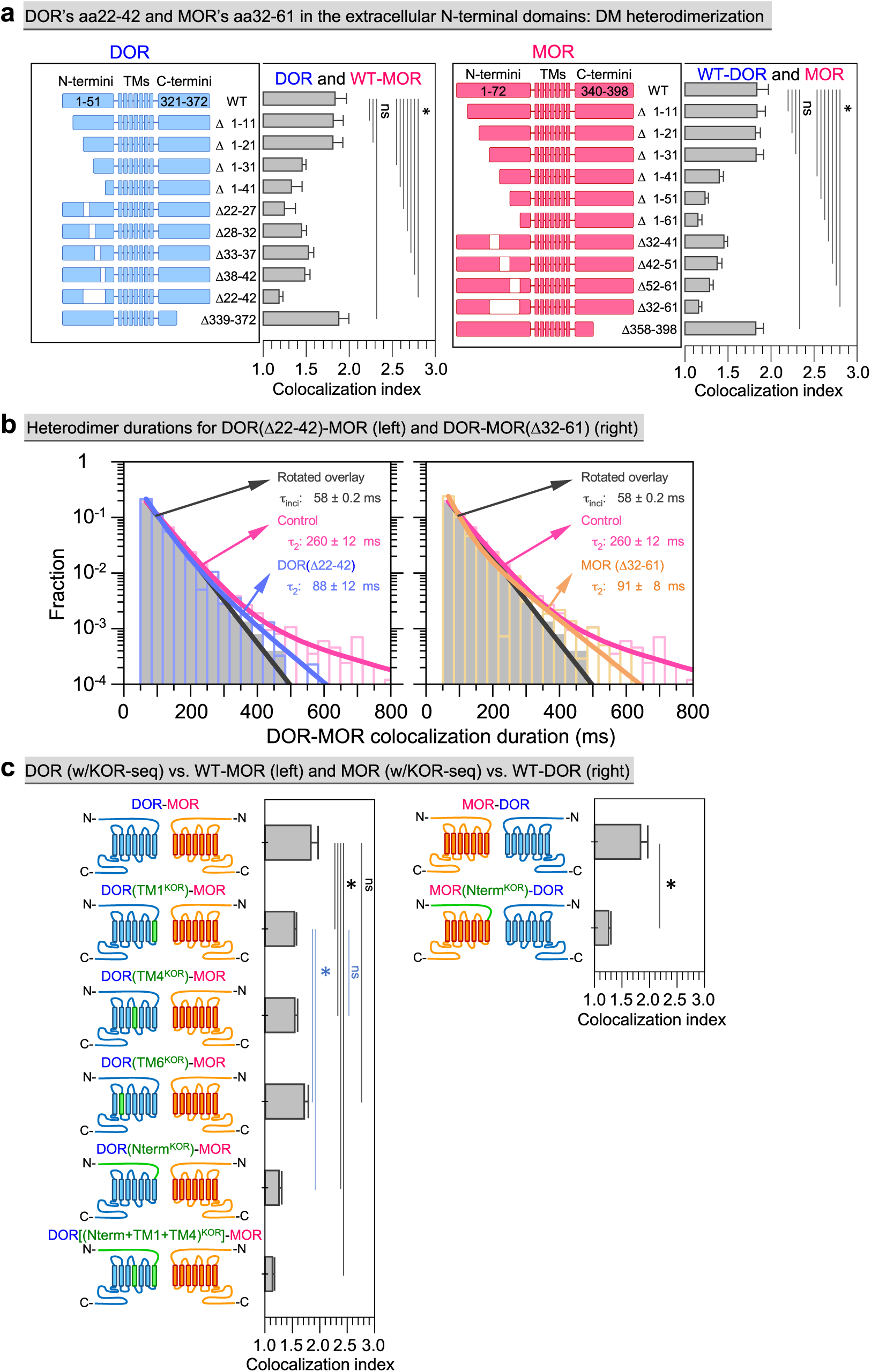
N-terminal domains of DOR’s aa22-42 and MOR’s aa32-61, as well as DOR’s TM1 and TM4 domains, are involved in DM heterodimerization. **a** Involvement of the extracellular N-terminal domains of DOR and MOR in DM heterodimerization, particularly that of DOR’s aa22-42 and MOR’s aa32-61. The figures show that mutants with various deletions in the N-terminal domains of DOR and MOR exhibit lower colocalization indexes with full-length (WT) MOR and DOR, respectively, compared to the colocalization index between full-length DOR and MOR. At the bottoms of both panels are the results for the C-terminal deletion mutants, indicating the lack of cytoplasmic C-terminal domain involvement in DM heterodimerization. **b** Histograms showing the colocalization *duration* distributions for the pair of WT-MOR and DOR(Δ22-42) (left, blue) and the pair of WT-DOR and MOR(Δ32-61) (right, magenta). Histograms for the control and rotated overlays of WT DOR-MOR images are the same as those shown in **Fig. 1d**. **c** The extracellular N-terminal domains of DOR and MOR are involved in DM heterodimerization (left and right panels, respectively), as are DOR’s TM1 and TM4 domains, but not the TM6 domain (left panel). (Left panel) Based on the finding that MK heterodimerization hardly occurs, we examined whether the DM colocalization index is reduced upon substitutions of DOR domains with the corresponding KOR domains, using WT-MOR as the partner. (Right panel) The DM colocalization index is reduced upon the substitution of MOR’s N-terminal domain with KOR’s N-terminal domain, which is not involved in DK heterodimerization (Fig. 2a), indicating the involvement of MOR’s N-terminal domain in DM heterodimerization.

### DOR’s EL3, TM1, and TM4, but not EL2, N- or C-terminal domain, contribute to DK heterodimerization

We first examined heterodimerization of wild-type (WT) KOR and N- or C-terminal deletion mutants (ΔN or ΔC mutants) of DOR (Fig. 2a, Top). We found that DOR’s N- and C-terminal domains are not involved in DK heterodimerization, and KOR’s ΔN or ΔC mutations are not involved in heterodimerization with WT-DOR (Fig. 2a, Bottom). These results indicate that DK heterodimerization involves neither the N- nor C-terminal domains of DOR and KOR.

Given the scarce occurrence of MK heterodimers (colocalization index = 1.15 ± 0.03), we tested whether MOR mutants with some domains substituted with DOR counterparts could heterodimerize with WT-KOR (Fig. 1e; see Fig. 2b for domain names and amino-acid numbers; results shown in Fig. 2c left). Substitution with DOR’s TM1 or TM4 domain in MOR (MOR[TM1^DOR^] or MOR[TM4^DOR^]) facilitated heterodimer formation with WT-KOR, qualitatively consistent with a previous result.^30^ However, replacing MOR’s EL2 or N-terminal domain (MOR[EL2^DOR^] or MOR[Nterm^DOR^]) did not induce heterodimerization with WT-KOR. Meanwhile, introducing DOR’s EL3 domain into MOR (MOR[EL3^DOR^]) resulted in heterodimerization with WT-KOR. These results indicate the involvement of DOR’s EL3 domain, alongside its TM1 and TM4 domains, in DK heterodimer formation.

The lifetime distribution for the MOR(EL3^DOR^)-KOR heterodimer revealed the appearance of the *1*_2_ component, which did not exist in that for the MK pairs (Supplementary Fig. 2). Although the *1*_2_ value is smaller than that for WT-DK heterodimers (164 ± 25 ms vs. 241 ± 9.7 ms), this result further suggests that DOR’s EL3 domain is involved in DK heterodimerization.

Experiments replacing various DOR domains with the MOR counterparts and examining heterodimerization with WT-KOR (based on the lack of MK heterodimerization) showed reduced heterodimerization for TM1, TM4, and EL3 domain replacements, but not for EL2 or N-terminal domain substitutions (Fig. 2c right). This further indicates that DOR’s EL3, TM1, and TM4 domains are integral to DK heterodimer formation.

### Soluble EL3-domain peptides of DOR and KOR both block DK dimerization

A variety of deletions (Fig. 2a) and substitutions (Fig. 2c) introduced into WT-DOR could alter its conformation, which might affect true interaction sites, leading to the reduction of DK heterodimerization. To study this aspect, we synthesized soluble peptides with the EL3 sequences of DOR and KOR (Fig. 1e). Addition of either peptide (1 µM, final concentration) to cells co-expressing DOR and KOR significantly reduced DK heterodimerization, unlike the peptide with the MOR EL3 sequence (Fig. 2d). This highlights the critical roles of the DOR and KOR EL3 domains in DK heterodimerization. Given that DOR’s EL2 domain and the N-terminal domains of both DOR and KOR are not involved in DK heterodimerization (Fig. 2a, c), these results indicate that the interaction between the DOR and KOR EL3 domains is crucial for DK heterodimerization, although that with KOR’s EL2 domain has not been ruled out.

Furthermore, we found that DOR’s TM1 and TM4 peptides expressed in the PM became colocalized with KOR, supporting their involvement in DK heterodimerization (Fig. 2b, e). These results are consistent with those shown in Fig. 2c and a previous observation,^31^ and indicate that the interactions between the DOR and KOR EL3s, and those of DOR’s TM1 and TM4 with KOR’s TM domains are involved in DK heterodimerization.

### DM dimerization is predominantly mediated by N-terminal domain interactions

To identify the amino-acid sequences responsible for DM heterodimerization, first, we systematically deleted partial sequences from the N- and C-terminal domains of DOR and MOR (Fig. 3a). We found that DOR’s N-terminal amino acids 22-42 (with the critical amino acids 22-27) and MOR’s N-terminal amino acids 32-61 play key roles in DM heterodimerization, likely by binding to each other (Fig. 3a). The key DOR sequence was further confirmed by replacing DOR’s amino acids 21-27 with an arbitrarily selected amino-acid sequence, rather than deleting the original sequence (Supplementary Fig. 3). Therefore, these sequences are likely to be the binding interfaces for DM heterodimerization. However, the colocalization index did not decrease to 1 (but to 1.15, as found for MK heterodimerization), probably due to the presence of other binding sites such as TM domains, as suggested previously (see below).^3, 43, 46^

The histograms of the colocalization *durations* for DORΔ22-42 with WT-MOR and MORΔ32-61 with WT-DOR both revealed only small fractions of the *1*_2_ components, with significantly reduced heterodimer lifetimes of 88 ± 12 ms and 91 ± 8 ms, respectively (Fig. 3b). These shorter lifetimes (≈90 ms), compared to the 260-ms lifetime of the WT DM heterodimers, might represent the lifetimes of DM heterodimers induced by the less-stable TM domain interactions.

We further explored the specific regions mediating DM heterodimerization by taking advantage of the fact that MK heterodimerization hardly occurs at the expression levels we employed. Thus, we examined WT-MOR interactions with DOR mutants with various DOR domains substituted with their KOR counterparts (Fig. 3c left). The colocalization index upon replacement of the N-terminal domain substantially decreased, consistent with the previous demonstration that the N-terminal domains of DOR and MOR are instrumental in DM heterodimer formation (Fig. 3a, b).

Substitution of DOR’s TM1 or TM4 with its KOR counterpart, but not TM6 (Fig. 1e), yielded moderate reductions in colocalization with WT-MOR, but not as much as the N-terminal domain substitution (with Nterm^KOR^), suggesting that DOR’s TM1 and TM4 play ancillary roles in DM heterodimer stability (Fig. 3c left). These results, but not that of TM6, are consistent with previous computational modeling predictions.^43, 46^ The DOR mutant bearing the three replacements of KOR’s N-terminal domain, TM1, and TM4 exhibited a lower colocalization index with WT-MOR, comparable to that for the MK hetero-pair (1.15).

Conversely, the replacement of MOR’s N-terminal domain with that from KOR (with Nterm^KOR^) greatly reduced the colocalization with DOR (Fig. 3c right). This result further supports the idea that the major interaction sites for DK dimerization do not involve the N-terminal domains, but the key interactions occur between the EL3 domains as well as TM domains in DOR and KOR (Fig. 2). Importantly, it further demonstrates that the predominant interactions for DM heterodimerization occur between the N-terminal domains of both DOR and MOR (Fig. 3a, b). The interactions of the N-terminal domains with the EL2 and EL3 domains would be far less important for DM heterodimerization than those between the DOR and MOR N-terminal domains, because the lack of either DOR residues 22-42 or MOR residues 32-61 is sufficient to significantly lower the colocalization index (Fig. 3a) and eliminate the second colocalization lifetime component (Fig. 3b).

### TM interactions enhance both homo- and hetero-dimerizations of ORs

The results described in the previous subsection further support the involvement of DOR’s TM1 and TM4 in both DM (Fig. 3c left) and DK (Fig. 2c, left and right, and e) heterodimerizations. Furthermore, MOR’s TM1 participates in both DM heterodimerization and MM homodimerization (Fig. 5b-e in companion paper; Fig. 5 of this paper). These results suggest that the TM interactions support both homo- and hetero-dimerizations of ORs; namely, the TM interactions are less specific. This finding is aligned with previous controversial reports: The conclusion we obtain from the reported results in the literature is that the same TM domains contribute to both homo- and hetero-dimerizations of ORs and other GPCRs to various extents.^43, 44, 46, 47, 48^

### Soluble N-terminal-domain peptides block DM heterodimerization

We then utilized the soluble peptides corresponding to the N-terminal sequences of both DOR and MOR implicated in DM heterodimerization, as we did for confirming the DK heterodimer binding sites (EL3 domains of both molecules; Fig. 2d). These soluble peptides from DOR and MOR are called Dpep(m-n)DM and Mpep(m-n)MD, respectively, where m and n indicate the amino acid numbers in the original OR sequences (see “Location in DOR” and “Location in MOR” in Fig. 4a, respectively). Among the various DpepDMs and MpepMDs, Dpep(20-42)DM and Mpep(32-61)MD most potently lowered the colocalization indexes to 1.2-1.3 (Fig. 4a), mirroring the effects observed with the deletion and replacement mutants (Fig. 3a, c).

**Fig. 4:**
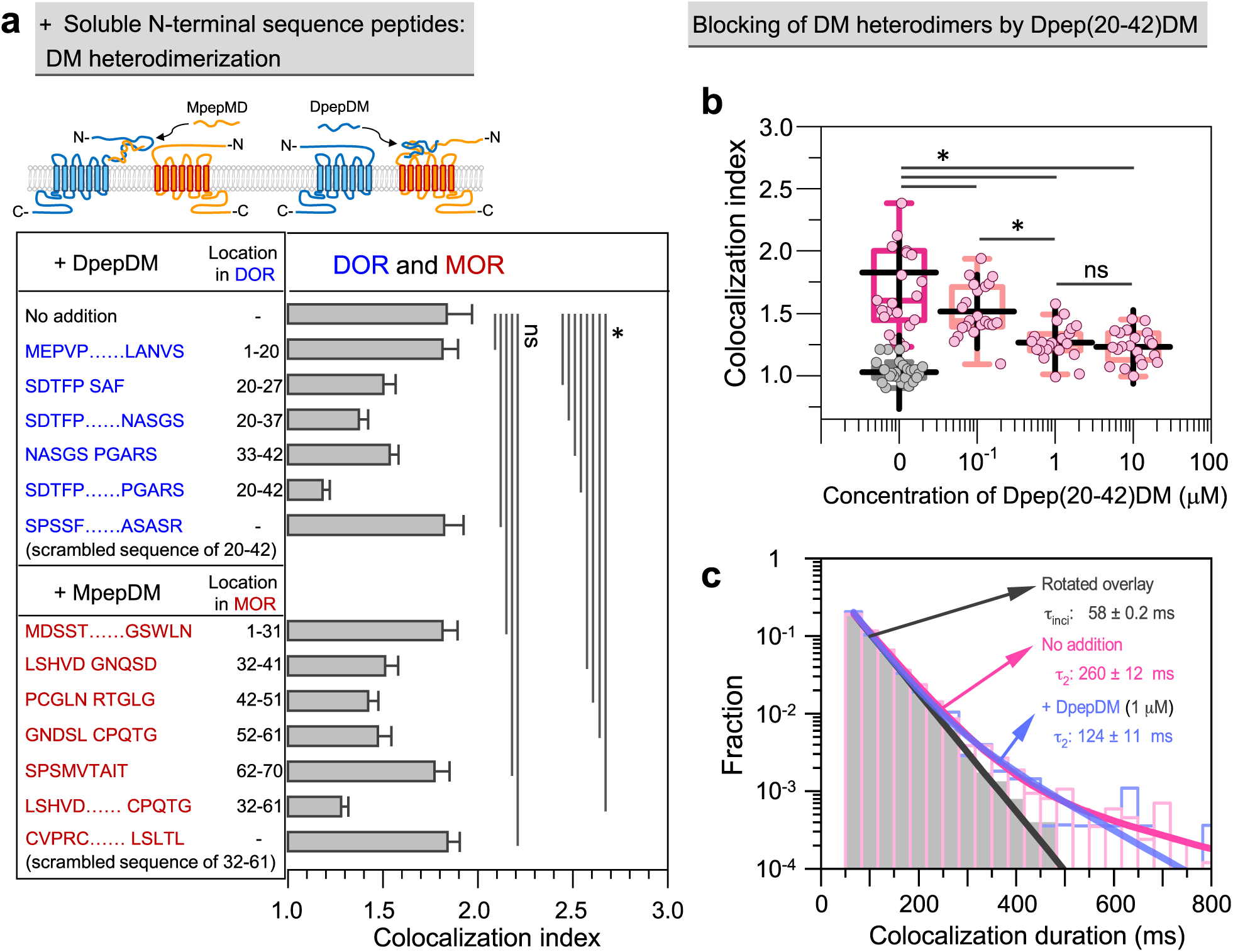
Soluble Dpep(20-42)DM and Mpep(32-61)DM peptides suppress DM heterodimerization. **a** Effect of the addition of 1 µM soluble peptides with the same amino-acid sequences as various parts of the N-terminal domains of DOR and MOR (DpepDM and MpepDM, respectively) on the DM colocalization index. **b** With an increase of the Dpep(20-42)DM concentration, the DM colocalization index decreased. The grey keys at 0 µM Dpep(20-42)DM represent the data points obtained by using rotated overlays. **c** Dpep(20-42)DM reduces the 1_2_ of DM colocalization (blue). Histograms for the control (no addition) and rotated overlays of WT DOR-MOR images are the same as those shown in **Fig. 1d**.

The colocalization index for DM heterodimers decreased with an increase in the Dpep(20-42)DM concentration (Fig. 4b), indicating that the affinity of the DM heterodimers is *K*_D_≈0.1 µM in three-dimensional space. This 3D concentration could not readily be converted to a two-dimensional molecular density, as discussed in the companion paper (main text explaining Fig. 5a-d). Briefly, in two-dimensional space, the dimerization efficiency is far greater, by a factor of 10^6^, than that in three-dimensional space.^49^ Therefore, the efficacy of the peptides in blocking OR dimerization, even at the 0.1 µM-order concentrations required for blocking DM heterodimerization, is deemed plausible given the nanomolar affinity range of OR ligands (1-10 nM).

**Fig. 5:**
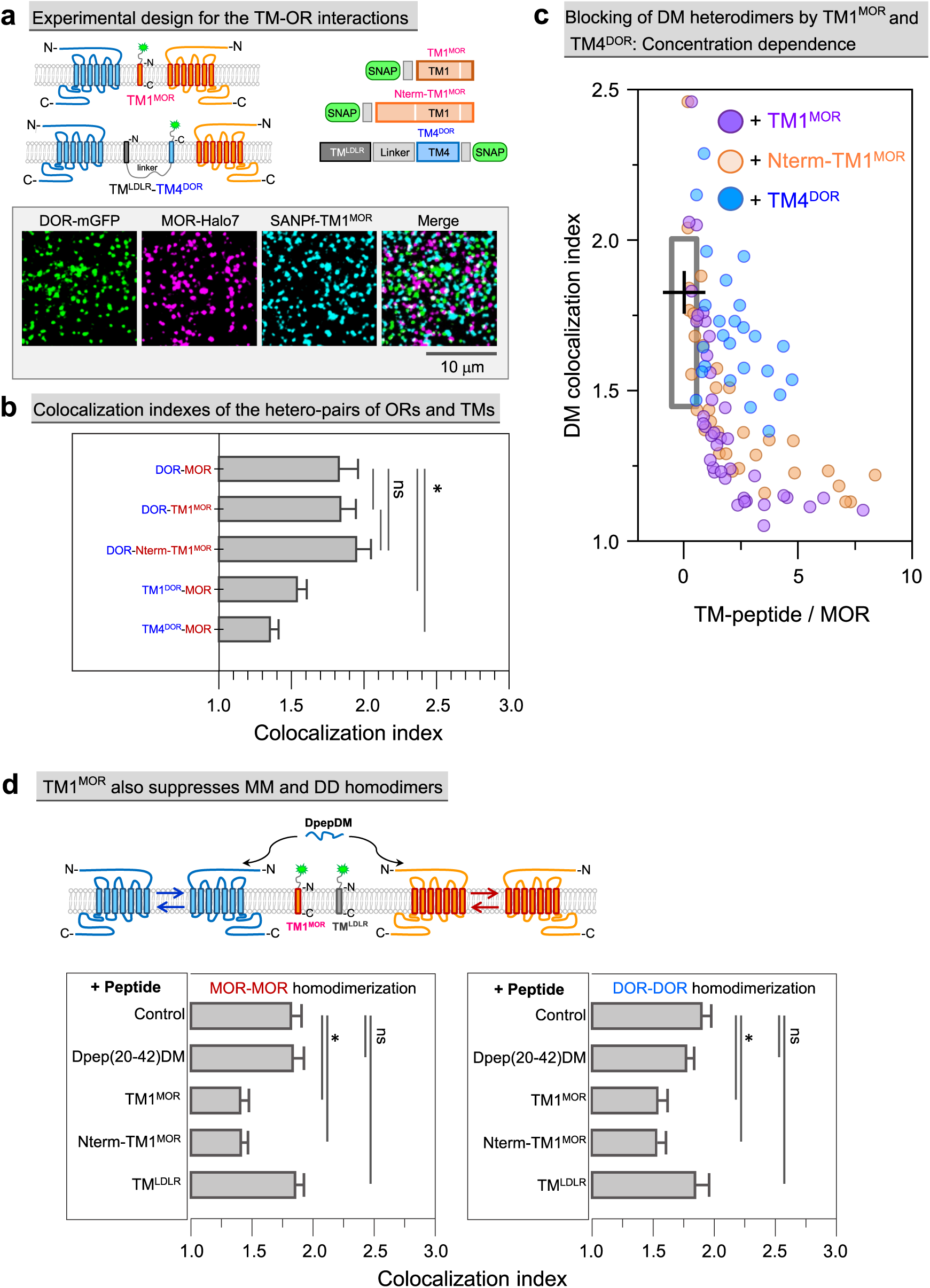
TM1MOR interacts with DOR and blocks DM heterodimers, and additionally inhibits MM and DD homodimers, whereas Dpep(20-42)DM only blocks DM heterodimer formation. **a** Experimental design for examining the interactions of TM1^MOR^, Nterm-TM1^MOR^, and TM4^DOR^ (Fig. 2b) with WT-ORs (top). For the correct orientation of TM4^DOR^ in the PM, it was linked to the transmembrane domain of LDL receptor (TM^LDLR^).^75^ Representative single-molecule images of DOR-mGFP (green), MOR-Halo (magenta), and SNAPf-TM1^MOR^ (cyan) co-expressed at similar levels of 0.5-1.0 fluorescent spots/µm^2^ (bottom). **b** TM1^MOR^ and Nterm-TM1^MOR^ form dimers with WT-DOR as efficiently as WT-MOR does. TM1^DOR^ and TM4^DOR^ form dimers with WT-MOR to lesser extents. Expression levels of all molecules were adjusted to 0.5-1.0 fluorescent spots/µm^2^. **c** Reduction of the DM colocalization index with an increase in the number densities of the TM peptides (TM1^MOR^, Nterm-TM1^MOR^, and TM4^DOR^) relative to that of MOR, expressed in the PM. DOR and MOR were both expressed at levels of 0.5-1.0 fluorescent spots/µm^2^. The effects of TM1^MOR^ and Nterm-TM1^MOR^ are stronger than that of TM4^DOR^. The rectangular box without the peptides (x = 0) indicates the result of the box plot without datapoints (to avoid excessive complexity in the plot). **d** TM1^MOR^ and Nterm-TM1^MOR^ reduce MM and DD homodimers (in addition to DM heterodimers), but 1 µM Dpep(20-42)DM does not. TM1^MOR^ and Nterm-TM1^MOR^ expression levels were ≈10x of MOR and DOR, following the results shown in panel **c**. TM^LDLR^ is a negative control, representing non-interacting TM domains.

Dpep(20-42)DM’s introduction substantially shortened the DM heterodimer lifetime from 260 ms to 124 ms (Fig. 4c), supporting that the amino acid sequence 20-42 in DOR is critical for DM heterodimer formation. The shortened lifetime might reflect heterodimer facilitation by the TM domains.

### The membrane-integral peptide TM1^MOR^ suppresses both DM heterodimerization and MM homodimerization

Next, we examined whether MOR’s TM1 (TM1^MOR^) is involved in DM heterodimerization, as previously described.^3, 43, 46^ In addition to TM1^MOR^ (amino acids 71-99), a second construct containing amino acids 53-70 before the TM1 domain (Nterm-TM1^MOR^) was used (see Fig. 2b for construct names and exact amino-acid ranges), following a previous report.^3^ When these peptides were co-expressed with DOR, both TM1^MOR^ and Nterm-TM1^MOR^ colocalized with DOR, like the full-length MOR (all molecules expressed at densities of 0.5-1.0 fluorescent spots/µm^2^; Fig. 5a, b). These TM1 constructs also suppressed DM heterodimerization in an expression level-dependent manner, indicating TM1^MOR^’s significant contribution to the DM interaction (Fig. 5c).

Notably, the addition of either Dpep(20-42)DM (also Mpep(32-61)MD) or TM1^MOR^ alone reduced the colocalization index to near-baseline levels (1.2-1.3) (Fig. 4a, b and 5c). These results suggest that DM heterodimerization might not be induced by simple bindings at two sites, but instead mediated by cooperative allosteric conformational changes of the extracellular N-terminal domains and TM domains in both OR molecules. In this case, our description of the dynamic dimer-monomer equilibrium should be considered as a simplified representation of the actual molecular events.

To identify the DOR’s TMs interacting with MOR, we examined the colocalization of WT-MOR with TM1^DOR^ and TM4^DOR^, which were previously suggested to interact with TM1^MOR^.^43, 46^ TM1^DOR^ and TM4^DOR^ moderately colocalized with MOR when expressed at similar levels (both at densities of 0.5-1.0 fluorescent spots/µm^2^; Fig. 5b). Furthermore, TM4^DOR^ suppressed DM heterodimerization in an expression level-dependent manner, but not as much as TM1^MOR^ (Fig. 5c), indicating its contribution to DM heterodimerization but also suggesting the presence of other significant binding sites, including TM1^DOR^ and other TM domains as well as the N-terminal domain.

Despite these findings, we emphasize the critical role played by the interactions of the N-terminal domains of DOR and MOR. This is because the TM interactions are not quite specific, as described. Namely, TM1^DOR^ and TM4^DOR^ are involved in both DM (Fig. 3c left) and DK (Fig. 2c, left and right, and e) heterodimerizations, and TM1^MOR^ is involved in both DM heterodimerization (Fig. 3c right; Fig. 5d as it blocks DD homodimerization) and MM homodimerization (Fig. 5d; also refer to Fig. 5b-e in companion paper). In striking contrast, Dpep(20-42)DM had no influence on either MM or DD homodimerization (Fig. 5d). A control TM-peptide, TM^LDLR^, with the sequence of the LDL receptor’s TM domain, exhibited no effect on homodimer blocking (Fig. 5d).

In conclusion, the N-terminal domains of DOR and MOR are specifically involved in DM heterodimerization, whereas the TM domains contribute but interact less specifically: TM1^MOR^ in MOR enhances both DM heterodimerization and MM homodimerization, and TM1^DOR^ and TM4^DOR^ in DOR participate in both DM and DK heterodimerization. Meanwhile, DM heterodimerization might involve allosteric interactions between the N-terminal and TM domains.

### M-agonists’ effect on DM heterodimerization

We examined the effects of MOR agonists (M-agonists), morphine and [D-Ala2, N-MePhe4, Gly-ol]-enkephalin (DAMGO), on DM heterodimerization (Fig. 6a-e). At an M-agonist concentration of 0.5 µM in the cell culture medium, MOR would be rapidly bound by morphine and DAMGO (denoted as M*^mor^ and M*^DAMGO^, respectively),^50^ and therefore we examined the M-agonist effect on DM heterodimerization within 5 min after the agonist addition, prior to any detectable internalization. As detailed in the companion paper, DAMGO enhanced MM *homo*dimerization and slightly elongated the MM dimer lifetime (Fig. 5c and d, respectively, in companion paper).

**Fig. 6:**
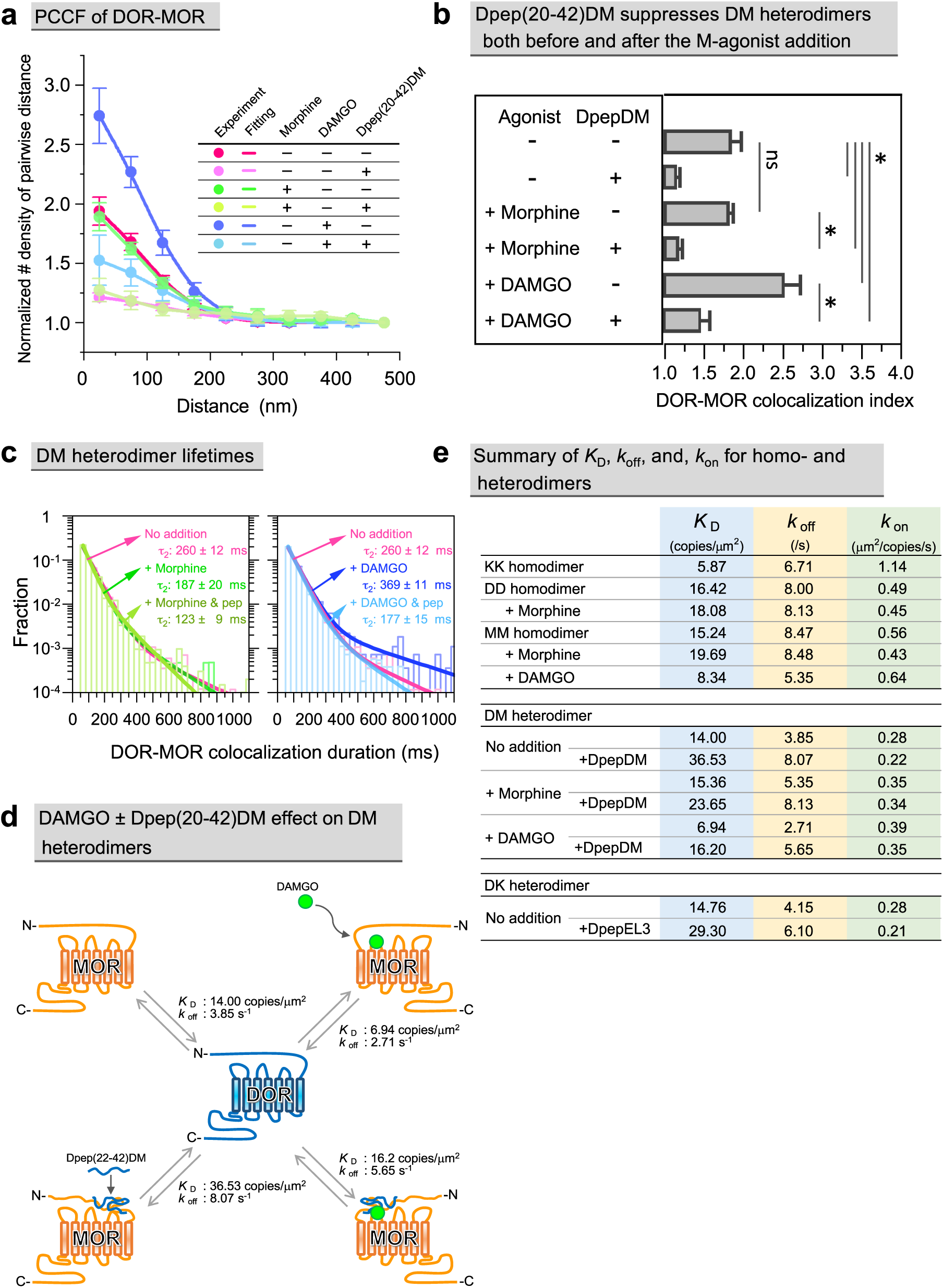
Effects of M-agonists, morphine and DAMGO, on DM heterodimerization, and of Dpep(20-42)DM on DM heterodimerization, in the presence of M-agonists. **a** Experimental PCCFs for DM heterodimerization in the presence and absence of 0.5 µM M-agonists and 1 µM Dpep(20-42)DM and their fitting curves using *K*_D_ for DM heterodimerization and σ (precisions of single-molecule localizations for the two probes and spatial precision for overlaying two-color images) as fitting parameters. MD-cells expressing both DOR and MOR at 0.5-1.0 fluorescent spots/µm^2^ were employed. **b** Colocalization indexes obtained from the PCCFs shown in **a**. Together with the results shown in **a**, the results indicate that morphine hardly affects DM heterodimerization whereas DMAGO enhances it. The Dpep(20-42)DM addition significantly reduces DM heterodimers under all conditions examined here. **c** Morphine shortens and DAMGO prolongs DM heterodimer lifetimes, whereas the further addition of Dpep(20-42)DM reduces the lifetimes, making them shorter than those for DM heterodimers without any treatment. **d** Schematic summary of *K*_D_ and *k*_off_ values for DM heterodimers in the presence and absence of DAMGO and DpepDM(20-42)DM. **e** Summary of *K*_D_, *k*_off_, and *k*_on_ values for OR homodimers and DM and DK heterodimers, including the cases in the presence and absence of morphine or DAMGO and DpepDM(20-42)DM (DpepEL3 in the case of DK heterodimers), wherever appropriate.

M*^mor^ exhibited almost the same colocalization index with DOR as non-ligated MOR, although the DM*^mor^ heterodimer lifetime was shorter (Fig. 6a-c). The effects of Dpep(20-42) on DM dimerization for M*^mor^ and non-ligated MOR were very similar (Fig. 6a, b, and e). M*^DAMGO^ exhibited a greater colocalization index with DOR (DM*^DAMGO^ heterodimers) with a prolonged heterodimer lifetime, compared to MOR in the basal state (Fig. 6a-e). The presence of Dpep(20-42)DM in the medium markedly reduced both the colocalization index and lifetime of the DM*^DAMGO^ heterodimers. The influence of the DOR agonist, 0.5 µM SNC-80, on MOR and SNC-80-bound DOR heterodimerization could not be determined due to the rapid internalization of MOR and DOR after SNC-80 addition, as reported in subsequent sections.

### Summary of *K*_D_, *k*_off_, and *k*_on_ values and expected percentages of OR protomers existing as homo- and hetero-dimers at various expression levels

The equilibrium constants *K*_D_s were deduced by fitting the experimental PCCF histograms for DM and DK heterodimers with theoretical models (Fig. 6a), using hetero-*K*_D_ and σ as fitting parameters (Eqs. 26 and 73-75 in Supplementary Note 2 in companion paper for homodimer). The derived *K*_D_, *k*_off_, and *k*_on_ values for KK, MM, and DD homodimers as well as DM and DK heterodimers, with or without M-agonists and Dpep(20-42)DM, are summarized in Fig. 6d and e (for the influence of expression levels on the evaluated *K*_D_ values with actual curve fitting results, see Supplementary Fig. 4a-d; for the SEM, see Supplementary Tables 2 and 3; all statistical data are summarized in Supplementary Table 1).

Based on the *K*_D_ values obtained here, the percentages of OR protomers existing as monomers, DD and MM homodimers, and DM and DK heterodimers in the presence and absence of morphine and DAMGO, at various number densities of ORs expressed in the PM, are calculated and shown in Fig. 7 and Supplementary Tables 2-4 for the cases where DOR and MOR expression levels are equalized. In particular, Supplementary Table 4 presents the fractions of MOR or DOR protomers existing as monomers, homodimers, and heterodimers. For the cases where expression levels are freely varied, see Fig. 7. Without knowing the *K*_D_ values for both homo- and hetero-dimerizations, such evaluations would have been impossible.

**Fig. 7.**
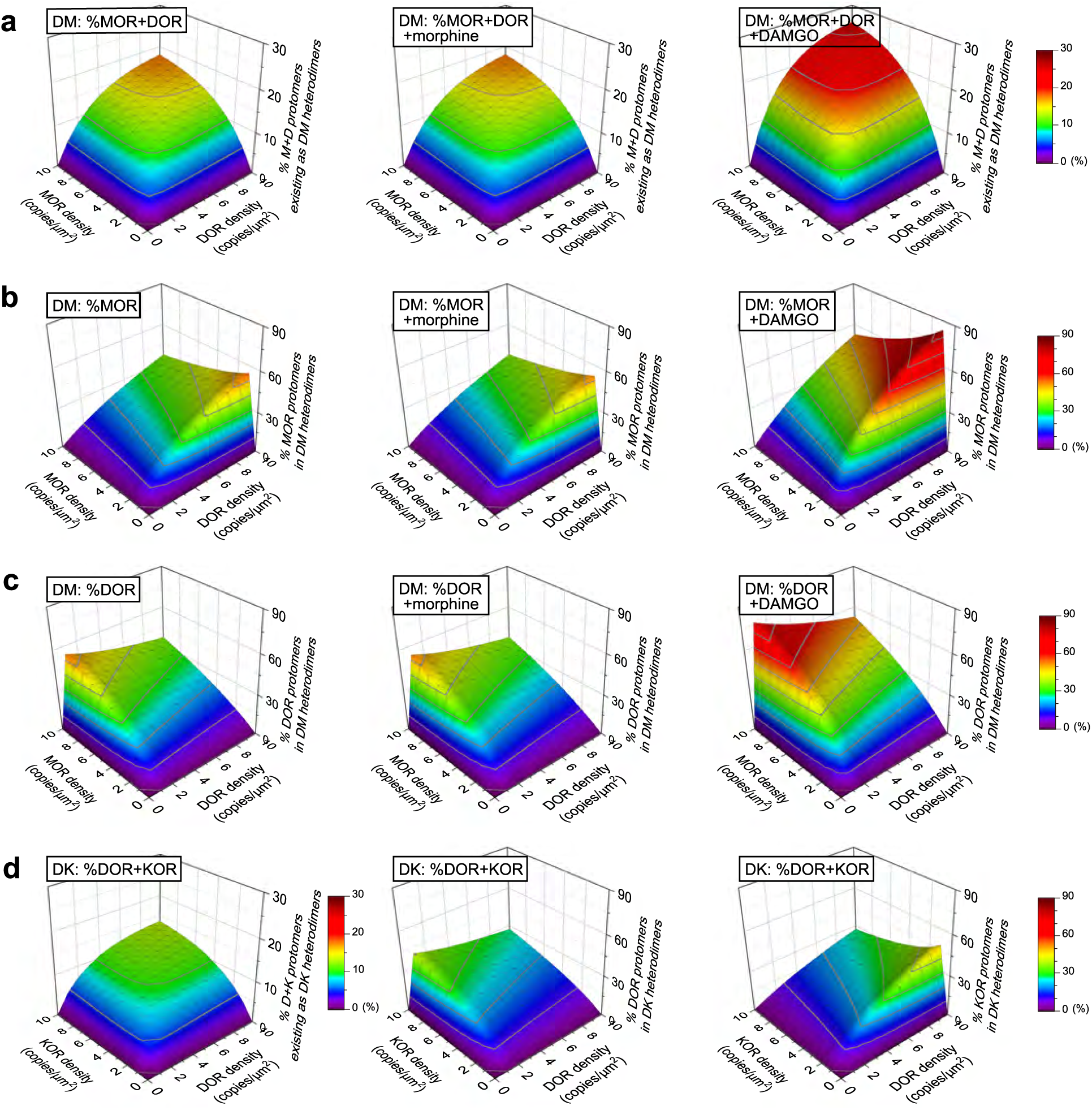
The percentages of OR protomers existing as OR heterodimers at various expression levels (z-axis), predicted from the *K*_D_ values for homo- and hetero-dimer dissociation equilibrium. In a-c, from left to right, no addition, + morphine, and + DAMGO. **a** Overall percentages of MOR and DOR protomers existing as DM heterodimers. **b** Percentages of MOR protomers existing as DM heterodimers. **c** Percentages of DOR protomers existing as DM heterodimers. **d** Overall percentages of DOR and KOR protomers existing as DK heterodimers (left), percentages of DOR protomers existing as DK heterodimers (middle), and percentages of KOR protomers existing as DK heterodimers (right).

### Virtually all OR molecules alternate between existing as homo- and hetero-dimers and monomers during short periods on orders of 1-10 s

Cautious interpretations are required for the small percentages of OR protomers, 4.2-21.2%, existing as homo- and hetero-dimers at any moment at low physiological expression conditions (1 copy/µm^2^ for each OR; see columns of expression levels of 1 copy/µm^2^ and 1+1 copies/µm^2^ in Supplementary Tables 2 and 3, respectively; including the presence of morphine and DAMGO, but not peptide dimer blockers). Based on these values, one might consider the functional influence of dimers to be small, but this would be incorrect. Note that the dimer lifetimes are in the range of 118-369 ms, indicating that these dimers fall apart quickly, but then the monomers will again form dimers with different (and sometimes the same) partner molecules (see Supplementary Fig. 2a and b in companion paper for an example). This process is continually repeated. Therefore, although at any time the percentages of protomers existing as dimers are limited, virtually all OR molecules rapidly interconvert among homodimer-, monomer-, and heterodimer-states and monomers during short periods on orders of 1-10 s (at physiological expression levels for both molecules) (Fig. 6e and Supplementary Table 3).

This indicates that ORs would perform their functions while undergoing rapid monomer-dimer interconversions. If the functionalities of monomers and dimers are different, this raises the possibility to fine-tune the OR functions by modulating monomer-dimer interconversions by natural agonists and synthetic drugs.

### Lack of morphine-induced internalization of MOR and DOR even in MD cells

Transient DM heterodimerization might affect agonist-induced intracellular signaling and internalizations of MOR and DOR, and thus analgesic efficacy and tolerance development (Fig. 8a). Therefore, we examined the effects of DM heterodimerization on downstream signals after M-agonist stimulation and the internalization of MOR and DOR before and after M- and D-agonist stimulation (0.5 µM), using three cell lines produced here: (1) M-cells, CHO-K1 cells stably expressing SNAPf-MOR at 0.5-1.0 fluorescent spots/µm^2^ after labeling with SNAP-Surface 549; (2) D-cells, CHO-K1 cells stably expressing SNAPf-DOR at 0.5-1.0 fluorescent spots/µm^2^ after labeling with SNAP-Surface 549; (3) MD-cells based on M-cells (or D-cells), transiently transfected with the cDNA encoding DOR-Halo (or MOR-Halo), which was labeled with SaraFluor 650T (cells exhibiting 0.5-1.0 fluorescent spots/µm^2^ were selected for observations). The two types of MD-cells (based on M-cells or D-cells) are biologically the same (thus, in the following, we simply call them MD-cells), apart from the experimental details (i.e., for the assay of MOR and DOR internalizations, SNAPf-MOR and SNAPf-DOR must be used, respectively, because our assay depends on the quenching of fluorescence signal from these molecules located on the cell surface using the membrane impermeable quencher; see below and the companion paper). Dpep(20-42)DM (1 µM) was extensively used to dissect the functions of DM heterodimers from those of D/M monomers and DD/MM homodimers.

**Fig. 8:**
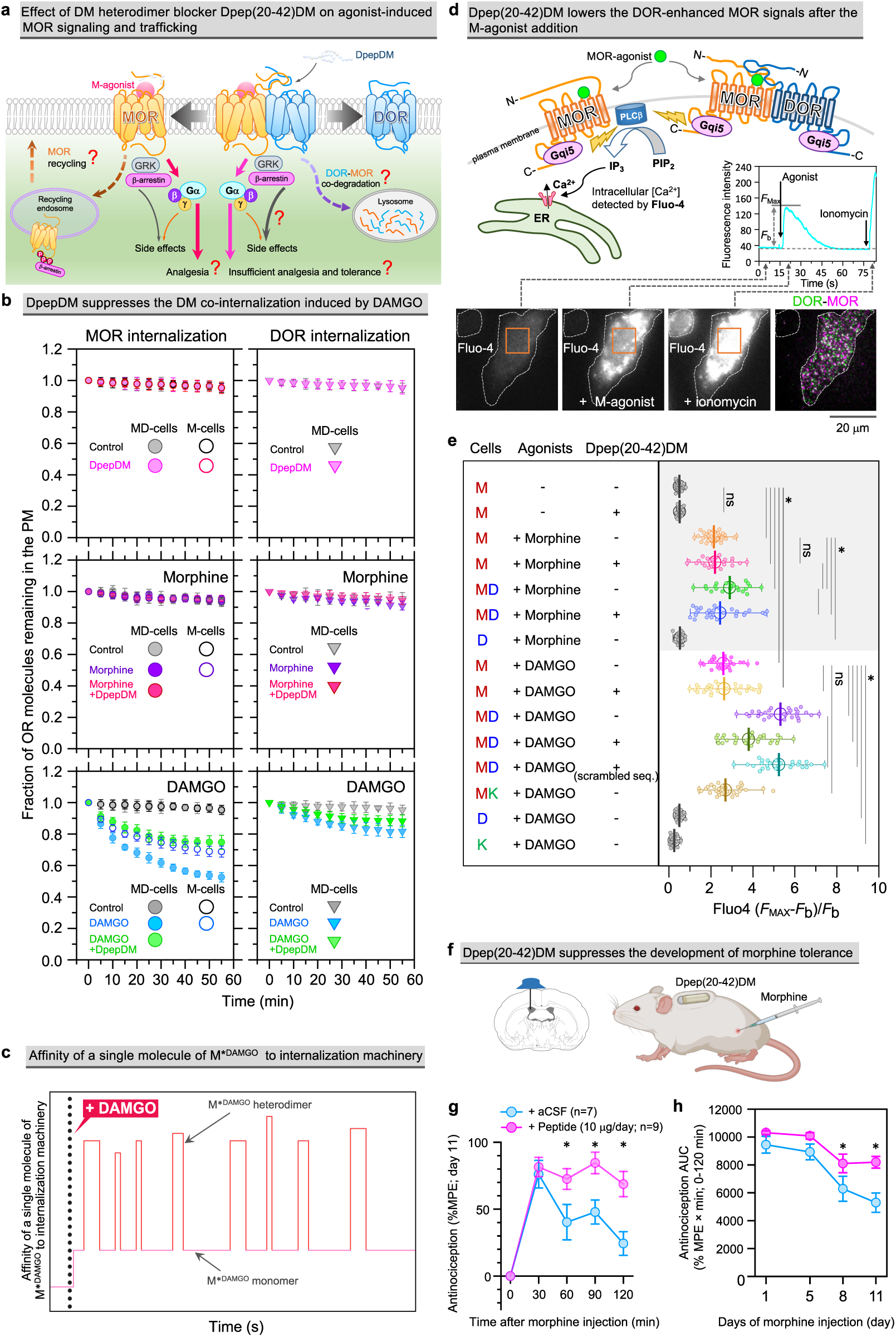
Dpep(20-42)DM helps to dissect the DM heterodimer and MOR monomer functions after M-agonist applications in cells, and enhances morphine analgesia and reduces tolerance development in mice. **a** Schematic figure showing that Dpep(20-42)DM reduces DM heterodimers, helping to dissect the DM heterodimer and MOR monomer functions, including signaling and internalization, upon M-agonist addition. **b** Time courses of MOR and DOR internalization (left and right columns, respectively) before and after the addition of 0.5 µM morphine or DAMGO (top, middle, and bottom boxes, respectively) in the presence and absence of 1 µM DpepDM. MOR internalization was examined in MD-cells and M-cells, whereas DOR internalization was observed only in MD-cells (MOR and DOR internalizations upon D-agonist addition were observed in MD-cells and D-cells, see Supplementary Fig. 6a). The fitting results for these time courses are not depicted here for clarity, but are shown in Supplementary Fig. 5. **c** Schematic illustration showing immensely enhanced efficiencies of binding to internalization machineries such as GRKs and β-arrestins and of DM*^DAMGO^ heterodimer internalization, occurring in the brief lifetimes of DM*^DAMGO^ heterodimers. M*^DAMGO^ monomers’ internalization efficiency is low, but due to the presence of many M*^DAMGO^ monomers, they are responsible for the internalization of half of the M*^DAMGO^ molecules in the PM. **d** (Top) Experimental design for observing Ca^2+^ mobilization after M-agonist addition. Ca^2+^ mobilization was monitored by Fluo-4 fluorescence intensity in M-, D-, and MD-cells. (Bottom) Fluo-4 images and single-molecule images of DOR and MOR. Ca^2+^ mobilization was parametrized by using [*F*_Max_-*F*_b_]/*F*_b_, where *F*_Max_ is the maximal Fluo-4 signal intensity within 75 s after stimulant addition and *F*_b_ is the baseline intensity (insert in Fig. d). **e** Ca^2+^ mobilization parametrized by using [*F*_Max_-*F*_b_]/*F*_b_ for Fluo-4 signals before and after the addition of morphine and DAMGO, in the presence and absence of Dpep(20-42)DM. **f** Schematic illustration showing the osmotic pump implantation for intracerebroventricular injection of Dpep(20-42)DM and subcutaneous morphine injection in mouse. **g** Tail flick test results on day 11, showing that morphine-induced analgesia was enhanced by the continuous administration of Dpep(20-42)DM (10 µg/day). The enhancing effect lasted for at least 120 min. **h** Tolerance to morphine was demonstrated by the loss of the analgesic response from day 8, reduced to ≈50% of the initial efficacy on day 11 in the control aCSF-treated mice (AUC = area under the antinociception vs. time curves shown in Supplementary Fig. 6b). In contrast, the antinociceptive effect of morphine in DpepDM-treated mice was maintained at ≈80% of the initial efficacy on days 8 and 11. All experiments were randomized and performed by a blinded researcher, who was then unblinded before statistical analysis.

The time courses of OR internalizations were monitored for 55 min by measuring the OR molecules remaining in the PM. The OR amounts in the PM were measured by TIRF microscopy, by evaluating the signal intensities of the OR molecules in the TIRF illumination range (in PM plus cytoplasm within ≈100 nm from the PM) before and after the addition of a membrane-impermeable fluorescence quencher, Mn(III) meso-tetra(4-sulfonatophenyl)porphine (Mn^3+^-TSP) (i.e., the intensity of the fluorescence signal from the cytoplasm in the TIRF illumination range), and then subtracting the latter from the former (see Fig. 6a and Supplementary Fig. 6 and related main text in the companion paper; also see Methods). Each time course obtained under various conditions could be fitted with a single exponential function plus a constant. The constant provided the OR fraction whose internalization is undetectable within 55 min, and the exponential decay constant provided the residency lifetime in the PM for the OR’s internalized fraction observed during 55 min (Fig. 8b and Supplementary Fig. 5; for summaries and statistical test results see Supplementary Table 5).

MOR internalization before and after morphine addition was hardly detectable in either M- or MD-cells (Fig. 8b, Left column, Top and middle; Supplementary Fig. 5, Top and third rows). These results indicate that the internalization of DM heterodimers is as slow as that of MOR monomers and homodimers and their internalization rates are not affected by morphine binding. This was further confirmed by the lack of Dpep(20-42)DM effects on M and M*^mor^ internalization (Fig. 8b, Left column, Top and middle; Supplementary Fig. 5, Top-fourth rows). DOR internalization was similarly unaffected by morphine addition in MD-cells (Fig. 8b, Right column, Top and middle; Supplementary Fig. 5, Top-fourth rows).

However, these observations require cautious interpretations due to the low fractions of M and M*^mor^ existing as DM and DM*^mor^ heterodimers, respectively, under the employed expression levels (≈5% as shown in Supplementary Table 3; see the column of 1+1 copies/µm^2^; i.e., the case approximating experimental observation conditions of 0.5-1.0 fluorescent spots/µm^2^ for both SNAPf-MOR and DOR-Halo), although, as described in the previous subsection, virtually all OR molecules would experience dimer periods within 1-10 s. In certain PM domains in some neurons, the MOR and/or DOR concentrations may far exceed those used in this study (e.g., MOR in primary cilia),^51^ and the morphine effect on the MM and/or DM dimer internalizations could be more pronounced.

### Highly-enhanced M*^DAMGO^ internalization during transient binding to DOR

In contrast to morphine, DAMGO administration resulted in pronounced internalization of DAMGO-bound MOR (M*^DAMGO^) in both M- and MD-cells (Fig. 8b, Left column, Bottom; Supplementary Fig. 5, Fifth and sixth rows; Supplementary Table 5). The results in M-cells are virtually the same as those reported in the companion paper (Fig. 6b, Middle column and Table 1), which show significant M*^DAMGO^ internalization during 35 min (55 min for the results shown here) that is not influenced by M*^DAMGO^M*^DAMGO^ homodimerization.

In MD-cells, the proportion of internalized M*^DAMGO^ is higher compared to that in M-cells (48% vs. 30%). This difference occurs despite the low fractions of M*^DAMGO^ existing as DM*^DAMGO^ heterodimers (9.1% at time 0) (Supplementary Table 3; see the 1+1 copies/µm^2^ column). The addition of Dpep(20-42)DM to MD-cells, which reduces heterodimeric M*^DAMGO^ from 9.1% to 4.3% (Supplementary Table 3), significantly reduced M*^DAMGO^ internalization from 48% to 26% (Fig. 8b and Supplementary Table 5). Therefore, we conclude that significant M*^DAMGO^ internalization occurs with DM*^DAMGO^ heterodimers in addition to M*^DAMGO^ monomers and homodimers. Although the co-internalization of MOR and DOR following DAMGO exposure has been documented previously,^3, 6, 52, 53, 54^ the critical insight obtained here is that DM*^DAMGO^ heterodimers, despite their very low fraction (9.1% of M*^DAMGO^ at low physiological expression levels of ≈1+1 copies/µm^2^ of MOR and DOR) and short lifetime (369 ± 11 ms; Fig. 6c, d), contribute to a significant proportion of M*^DAMGO^ internalization.

These observations suggest an unanticipated regulatory mechanism for M*^DAMGO^ internalization. Namely, at the moment of DM*^DAMGO^ heterodimer formation, its ability to bind to cellular internalization machineries, such as GRKs and β-arrestin 2,^55^ might suddenly rise dramatically, and then the binding affinity would rapidly decline due to DM*^DAMGO^ heterodimer dissociation, which would occur within a fraction of a second (1/*k*_off_ = 0.37 s; Supplementary Table 3). Hence, the cycle of DM*^DAMGO^ heterodimerization and dissociation (which occurs every few seconds or so) would function analogously to a car driver adjusting the speed by frequently and briefly pushing on and releasing the gas pedal, to regulate the M*^DAMGO^ internalization rate (Fig. 8c). Although the enhancement of internalization might occur only during the brief periods when DM*^DAMGO^ heterodimers are formed (Fig. 8c), the total frequency of heterodimerization across the PM ensures that the overall internalization of M*^DAMGO^ as DM*^DAMGO^ heterodimers is significant. This correlates with the area under the curve consisting of many short pulses, depicted in Fig. 8c.^56, 57, 58^ Thus, the overall internalization rate of M*^DAMGO^ could be modulated by the frequencies and durations of transient DM*^DAMGO^ heterodimerization events.

M*^DAMGO^ monomer/homodimer internalization would occur much less efficiently at the level of individual molecules, but since larger fractions of M*^DAMGO^ exist as monomers at any moment, the overall contributions of M*^DAMGO^ monomers/homodimers to M*^DAMGO^ internalization are quite significant (Fig. 8c). If the expression levels of MOR and DOR are elevated in certain cell types, then M*^DAMGO^ internalization would predominantly occur through DM*^DAMGO^ heterodimers.

We also examined the effects of the DOR-specific agonist SNC-80 on receptor internalization.^59^ The treatment led to extensive internalization of SNC-80-bound DOR (D*^SNC^) in both D- and MD-cells, with similar extents and rates (Supplementary Fig. 6a, middle and right columns; Supplementary Table 5), indicating that DOR*^SNC^ is internalized without the influence of heterodimerization with MOR. Meanwhile, SNC-80 induced significant MOR internalization in MD-cells, which was reduced by the further addition of Dpep(20-42)DM (Supplementary Fig. 6a, left column; Supplementary Table 5), indicating that MOR is internalized when it forms heterodimers with D*^SNC^ in MD-cells.

### Transient DM*^Ago^ heterodimers produce strong short pulse-like signals

We examined the cytoplasmic signals triggered by M-agonists, morphine and DAMGO, by monitoring the intracellular Ca^2+^ mobilization in cells stably expressing Gqi5 and MOR, which were further transiently transfected to express DOR or KOR (called M-, MD-, and MK-cells in this subsection; Fig. 8d; cf Supplementary Fig. 1a and Methods as well as the caption to Supplementary Fig. 1a in the companion paper). All ORs were expressed at the same low physiological expression levels employed for single-molecule detection of dimers (0.5-1.0 fluorescent spots/μm^2^ for each OR).

In M-cells, Ca^2+^ mobilization occurred at comparable intensities upon morphine and DAMGO treatments, without any effect of Dpep(20-42)DM (in the companion paper, which also reports Ca^2+^ mobilization in M-cells, we only used DAMGO as an MOR agonist). No response was evident in D-cells or K-cells (only DAMGO was examined for K-cells). In MD-cells, augmentations of the Ca^2+^ signal compared to that in M-cells were apparent for both morphine and DAMGO (Ca^2+^ signal increase in MD-cells was greater with DAMGO than morphine), which were mitigated by Dpep(20-42)DM. These observations indicate that the overall M*^Ago^ downstream signals (M*^Ago^ denotes agonist-bound MOR, collectively indicating M*^mor^ and M*^DAMGO^) are greater when DOR is co-expressed. This result is consistent with a previous report indicating that morphine and DAMGO activate MOR with greater efficacy when DOR is co-expressed over MOR alone, but without explicitly estimating the percentages of MOR existing as DM heterodimers.^50^

We found very small fractions of M*^Ago^ molecules existing as DM*^Ago^ heterodimers (4.9% for DM*^mor^ and 9.1% for DM*^DAMGO^ at low experimental expression levels of ≈1+1 copies/µm^2^ in Supplementary Table 3) for very brief durations (369 ± 11 ms for DM*^DAMGO^ and 187 ± 20 ms for DM*^mor^; Fig. 6c). In the case of DAMGO, M*^DAMGO^ molecules existing as a *homo*dimer induce the same signal level as M*^DAMGO^ existing as a monomer (since the presence of FAM-Mpep-TAT or mGFP-Mpep does not affect the Ca^2+^ signal level; Figure 6F in companion paper). These results suggest that the signal production rate (signal induced per unit time) of each M*^Ago^ molecule in a DM*^Ago^ heterodimer must be substantially larger than that of each monomeric or homodimeric M*^Ago^ molecule. This enhanced signaling rate may be induced by conformational changes of M*^Ago^ upon DOR binding and/or those of DOR upon M*^Ago^ binding, which might make a non-ligated DOR signal possible.

The enhanced signal from the M*^Ago^ molecules in very limited populations of DM*^Ago^ heterodimers existing for short periods (369 ms) suggests a fundamentally important signaling mechanism by DM*^Ago^ heterodimers, for the same reasons as the enhanced internalization upon DM*^DAMGO^ heterodimerization (previous subsection). Namely, at the moment of DM*^Ago^ heterodimer formation, the probability that the M*^Ago^ molecule and possibly the M*^Ago^-bound DOR molecule bind to and activate Gqi5 or Gαi, consequently triggering downstream signaling cascades, might rise steeply, but this activation is short-lived because the DM*^Ago^ heterodimer dissociates quickly within fractions of a second (1/*k*_off_ = 0.19 s and 0.37 s for DM*^mor^ and DM*^DAMGO^, respectively; Supplementary Table 3). Hence, a DM*^Ago^ heterodimer will create brief, pulse-like signals, and thus the signal in the entire cell is produced as the sum of the brief, stronger pulse-like signals from individual DM*^Ago^ heterodimers plus the weaker signals from M*^Ago^ monomers (Fig. 8c).^56, 57, 58^ As we proposed previously,^56, 57, 58^ the regulation of the signal time courses of an entire cell is more readily achieved when signaling entities (individual signaling molecules and molecular complexes) emit short pulse-like signals as opposed to prolonged ones: only simple summation (aggregation) of brief signals suffices to generate the overall cellular signal response, while if individual signals are prolonged, complex integration rather than simple summation must be performed to produce the required cellular signal time courses.^58^ In short, the cellular signal intensities might be regulated by adjusting the frequencies of the short pulse-like formation of DM*^Ago^ heterodimers. In addition, in cells with elevated but still physiological OR expression levels, the signals emanating from DM*^Ago^ heterodimers could be quite substantial, governing the cellular signal.

### Dpep(20-42) reduces morphine tolerance in mice

As described in the previous subsection, Dpep(20-42)DM moderately reduces the M*^Ago^-induced signal. This effect not only has implications at the cellular level but also might extend to broader physiological contexts, including nervous tissues and systemic functions. Therefore, we examined whether the soluble DM heterodimer blocker peptide Dpep(20-42)DM can modulate morphine-mediated analgesia in murine models. The peptide, dissolved in artificial cerebrospinal fluid (aCSF), was administered directly into the cerebral ventricles at a dose of 10 µg/day continuously for two days prior to the commencement of daily subcutaneous morphine injections (10 mg/kg/injection; one injection/day) (Fig. 8f). We utilized the tail-flick test to evaluate the analgesic efficacy of morphine and track the progression of antinociceptive tolerance.

Our results revealed that the analgesic potency of morphine was enhanced by Dpep(20-42)DM, which was especially evident between 60 and 120 minutes after morphine application over the course of the study (days 1, 5, 8, and 11; Supplementary Fig. 6b). Notably, on day 11, the disparity in analgesic effects between the control aCSF and the peptide groups reached ≈3 fold at the 120-minute mark following morphine administration (Fig. 8g).

Furthermore, the administration of Dpep(20-42)DM reduced the development of morphine tolerance. By day 8 after the morphine treatment was initiated, a noticeable reduction in morphine’s analgesic effect was observed in the control group. In contrast, mice treated with the peptide retained ≈80% of the initial antinociceptive efficacy, which was sustained from days 8 to 11 (Fig. 8h). These results imply that the inhibition of DM* heterodimerization by Dpep(20-42)DM diminishes the development of antinociceptive tolerance to morphine, suggesting the involvement of crucial neuronal populations within the pain pathway that co-express MOR and DOR, corroborating prior studies.^3, 18, 21, 24^ Suppression of DM* heterodimers by Dpep(20-42)DM also diminishes the cellular signal levels (Fig. 8e). Accordingly, maintaining proper signal levels in cells might be important for sustained morphine analgesia, which will be discussed further.

## Discussion

The significance of OR heterodimerization in pain therapy and neuropsychiatric disorders is garnering increasing attention.^7, 8, 9, 10^ However, our understanding of OR heterodimers and their clinical implications is limited due to the paucity of fundamental knowledge on heterodimer formation dynamics and mechanisms and the lack of specific blockers. Using our advanced single-molecule imaging and analysis methods,^1^ we obtained the fundamental parameters *K*_D_, *k*_off_, and *k*_on_, describing OR homo- and hetero-dimerizations. Crucially, these parameters are independent of OR expression levels, and thus we firmly established the formation of transient DM and DK heterodimers, alongside KK, MM, and DD homodimers, in the PM of cells expressing these molecules. These parameters were determined in the presence and absence of M-agonists and Dpep(20-42)DM (Fig. 6e and Supplementary Tables 2-3), and the percentages of protomers existing as DM and DK heterodimers at various expression levels were calculated from the *K*_D_ values (Supplementary Tables 3-4 and Supplementary Fig. 5), which will be useful for future studies to reveal the biological and pharmacological consequences of OR homo- and hetero-dimers. We hardly detected transient OR trimers and tetramers, indicating that at low physiological expression levels, the percentage of OR protomers existing as oligomers greater than dimers would be limited.

This study first documented the transient nature of OR heterodimers, revealing lifetimes of DM and DK heterodimers of approximately 250 ms, roughly twice as long as those of homodimers (120 ms for DD and MM and 180 ms for KK). Previous single-molecule imaging studies likely missed the OR dimers due to their scarce presence at the low expression levels typically employed in these studies (≈1 copy/µm^2^) and their brief lifetimes.^60, 61, 62^ Indeed, experiments performed at higher protein densities using bioluminescence resonance energy transfer (BRET), fluorescence resonance energy transfer (FRET), and biochemical assays detected the dimers, although these methods could not elucidate the dynamic nature of OR dimerization.^44, 61, 63, 64, 65^ Interestingly, pull-down assays, which require minute-order procedures, detected homo- and hetero-dimers despite their 100-ms-order lifetimes,^18, 66^ probably due to the rapid reassociation kinetics.

The interaction domains critical for heterodimerization have been identified as both the N-terminal domains for DM and both the EL3 domains for DK heterodimers (Fig. 2-4). While TM domains contribute to dimerization, they are less specific and primarily augment overall affinity. For example, TM1^MOR^ is involved in MM *homo*dimerization by binding to another MOR (perhaps mainly the TM1 domain), in addition to DM *hetero*dimerization by binding to TM1^DOR^, TM4^DOR^, and other TMs of DOR (Fig. 5b-d). The nature of lower-specificity TM interactions is also found in the literature. The involvement of TM domains in DM heterodimerization^3, 46^ and DK heterodimerization was previously proposed,^31^ whereas the same TM domains such as TM4^DOR^ and TM1^MOR^ were also proposed to be involved in DD^67^ and MM^48, 68^ homodimerizations. As described in the companion paper, specific C-terminal interactions are critical for the formation of KK, MM, and DD homodimers. The TM interactions with lower specificities were previously proposed as the rolling dimer interface model, in which multiple TM interaction sites co-exist and interconvert.^44^

Since the addition of either Dpep(20-42)DM (also Mpep(32-61)MD) or TM1^MOR^ alone reduced the colocalization index for DM heterodimers to near-baseline levels (1.2-1.3) (Fig. 4a, b, and 5c), we suggest that DM heterodimerization might also involve cooperative allosteric conformational changes between the N-terminal domains and TM domains in DOR and MOR, which might affect their functions.^69^ In addition, although the binding of nucleotide-free Gα to GPCR was proposed to affect GPCR conformations,^70^ we did not consider this effect on OR dimerization. The possible involvement of these processes suggests the complexity of OR dimerization, which we have simplified using first-order kinetics to aid our understanding at this stage of research.

Our findings also suggest that while TM peptides from ORs will not serve as specific blockers for OR homo- and hetero-dimerizations (Fig. 5c and d),^3^ soluble peptides with sequences mirroring the extracellular heterodimer formation domains (EL3^DOR^ and EL3^KOR^ peptides for DK heterodimers and DpepDM and MpepMD for DM heterodimers) can specifically reduce DM and DK *hetero*dimers (Fig. 2d, 4, and 5d).

We extensively utilized the Dpep(20-42)DM peptide to dissect the functions of M*^Ago^ monomers/homodimers and DM*^Ago^ heterodimers after the application of M-agonists, morphine and DAMGO (where M*^Ago^ collectively denotes agonist-bound MOR). DM*^Ago^ dimerization was enhanced by DAMGO but not morphine, and reduced by Dpep(20-42)DM (Fig. 6). Morphine does not induce the internalization of M*^mor^ or DOR even in MD-cells, where DM*^mor^ heterodimerization is induced. M*^DAMGO^ monomers and M*^DAMGO^M*^DAMGO^ homodimers are internalized in similar manners (Fig. 6b in the companion paper), and DM*^DAMGO^ heterodimerization further enhanced M*^DAMGO^ internalization (Fig. 8b and c). DM*^Ago^ heterodimerization also enhanced the downstream Ca^2+^ signal (Fig. 8e). Monomers of a prototypical GPCR, the β2-adrenergic receptor, were previously found to efficiently activate a stimulatory heterotrimeric G protein, Gs,^71^ but our results indicate that certain GPCRs, including MOR, might acquire enhanced activities upon heterodimer formation.

In our study, conducted under the low physiological receptor expression levels (≈1+1 copies/µm^2^), only small fractions (≤9.1%) of M*^Ago^ will exist as DM*^Ago^ heterodimers at any moment (Supplementary Table 3). Combining this with the short 369-ms lifetime of DM*^Ago^ heterodimers, we advanced an argument that each DM*^Ago^ heterodimer will create a brief, intense pulse-like signal, contrasted with the weaker signals from M*^Ago^ monomers, and the overall cellular signal is produced as the sum of these signals (Fig. 8c, e).^56, 57, 58^ Note that all M*^Ago^ molecules form DM*^Ago^ heterodimers every 1-10 s even under lower normal physiological expression levels (Fig. 6e and Supplementary Table 3). Enhanced internalization of M*^DAMGO^ might also follow a similar enhancement pattern. These results were observed under expression levels far lower than those typically used in bulk optical and biochemical assays (by factors of 10-1000),^63, 64, 65^ suggesting a unique signaling mechanism in native physiological conditions.

In specific neural tissue domains or certain cell types where OR expression levels are higher,^51^ DM*^Ago^ heterodimers could play more dominant role in cellular responses, dictated by the frequency of their transient active phases. Building upon this hypothesis, and recognizing the possibility that small quantities of DM*^Ago^ heterodimers might induce critical signals in key neurons, we explored the *in vivo* effects of Dpep(20-42)DM administered into murine cerebral ventricles to assess its impact on morphine tolerance, unlike a previous study using systemically applied Nterm-TM1^MOR^-TAT,^3^ which would non-specifically block multiple dimer forms (Fig. 5). We found that cerebral ventricle-administered Dpep(20-42)DM significantly curtails the development of long-term tolerance (Fig. 8f-h). However, directly comparing the effectiveness of Dpep(20-42)DM to Nterm-TM1^MOR^-TAT is challenging due to differing administration methods and the inherent specificity of Dpep(20-42)DM, which likely contributes to its reduced side effects.

Our *in vivo* findings align with previous findings that a genetic or pharmacological blockade of DOR can enhance MOR-agonist induced analgesia, highlighting the role of DM heterodimers in regulating MOR-mediated pain pathways^4, 18, 21, 24, 72, 73, 74^ The newly identified specific extramembrane domains identified for DM heterodimer-blocking present novel targets for modulating OR function, offering strategic avenues for the development of opioid therapies with reduced tolerance and dependence.

These *in vivo* results should be compared with our findings in cultured cells, where Dpep(20-42)DM, facilitating DM*^mor^ dissociation into monomers, reduces the Ca^2+^ signal from Gqi5 (Fig. 8e). These apparently contrasting findings made *in vivo* and *in vitro* suggest that finely tuning cellular signals upon morphine application is crucial for preventing long-term tolerance in mice. Agents that modulate the levels of DM*^mor^ heterodimers in the PM, like Dpep(20-42)DM, could offer more precise control over drug effects than simply varying morphine dosage.

## Methods

### cDNA construction

See the companion paper under the same title.^1^

### Mice

Male C57BL/6N mice (30-40 g) were purchased from CLEA Japan, Inc. and maintained on a 12-h light/dark cycle with rodent chow and water available ad libitum. They were housed in groups of four until testing. Animal studies were performed according to protocols approved by the Nagasaki University Animal Care and Use Committee (Approved number: 2002181596).

### Cell culture, transfection, and microscope observations

See the companion paper under the same title.^1^

### Fluorescence labeling of ORs

The SNAPf-tag-protein fused to ORs was fluorescently labeled with the SNAP-Surface 549 ligand, basically as described in the companion paper.^1^ The Halo7-tag protein fused to the intracellular C-terminus of the OR (Supplementary Fig. 1a) was covalently conjugated with SaraFluor650T-Halo-ligand (Goryo Kayaku), by incubating the cells with 100 nM Halo ligand. The SNAPf-OR and OR-Halo7 proteins were simultaneously labeled in the growth medium, at 37°C in the CO_2_ incubator for 30 min. The cells were washed three times with fresh medium (5-min incubation each time), and then the Ham’s F12 observation medium was added.

The labeling efficiency was determined by using the SNAPf-CD47-mGFP and mGFP-CD47-Halo7 proteins expressed in the PM (Supplementary Fig. 1c). CD47 is a monomeric 5-pass transmembrane protein expressed in the PM.^76^ We could not use ORs for this purpose because they form transient dimers, and thus employed monomeric CD47. For further details, refer to Methods and Supplementary Fig. 1c-e in the companion paper.^1^

The fluorescence labeling efficiencies under our normal conditions were 60% and 88% for SNAP-Surface 549 and SaraFluor650T, respectively (Supplementary Fig. 1c). By controlling the expression levels and using these labeling conditions, the spot numbers of SNAPf-OR and OR-Halo7 proteins were approximately equalized (0.75 ± 0.25 spots/µm^2^ for each color; i.e., a total spot number density of 1.5 ± 0.5 spots/µm^2^).

### Single fluorescent-molecule imaging

Fluorescently labeled ORs expressed in the bottom PM (the PM facing the coverslip) at fluorescent-spot number densities of 0.75 ± 0.25 spots/µm^2^ for each color (a total spot number density of 1.5 ± 0.5 spots/µm^2^) were observed at the level of single molecules at 37°C, using a home-built objective lens-type TIRF microscope constructed on an inverted microscope (Olympus IX-83) (see the companion paper for details).^77, 78, 79^ ORs tagged with fluorescent probes were excited with TIR illumination using the following power densities: SNAP-Surface 549 at 561 nm (Coherent OBIS 561-100 LS) at 0.35 µW/µm^2^; and SaraFluor650T at 642 nm (Omicron LuxXPlus 640-140) at 0.50 µW/µm^2^. Simultaneous two-color single-molecule imaging and analysis were conducted as described in the companion paper.^79^

### Evaluating colocalization durations

See the companion paper under the same title. The trackable duration lifetime (*1*_track_) for SNAP-Surface 549 bound to SNAPf-MOR was 16.3 ± 1.2 s (n = 500; the same as those described in the companion paper) and that for SaraFluor650T-Halo-ligand bound to MOR-Halo was 12.5 ± 0.8 s (n = 500) (expressed in CHO-K1 cells) (Supplementary Fig. 1f).

### Evaluating the colocalization index

See the companion paper under the same title.^1^

### Monte-Carlo simulations

See the companion paper under the same title.^1^

### Cell treatments with agonists and peptides

M-agonists, DAMGO (Sigma-Aldrich) and morphine (Takeda), and D-agonist SNC-80 (Sigma-Aldrich) were applied to the cells as described in the companion paper. DpepDM and MpepMD peptides, custom-synthesized by Cosmo-Bio, are soluble peptides, and were dissolved in the observation medium and added to the cells in the observation medium at a selected final peptide concentration at 37°C. Since their target sites are located on the cell surface, the complex procedures employed for incorporating the FAM-Xpep-TATs inside the cells, as described in the companion paper, were unnecessary.

### Ca^2+^ mobilization assay

The Ca^2+^ mobilization assay was performed in basically the same way as described in the companion paper. However, when we observed Ca^2+^ responses using the cells expressing SNAPf-MOR, we employed the CHO-K1 cells stably expressing both Gqi5 and SNAPf-MOR (at number densities of 0.75 ± 0.25 spots/µm^2^ when labeled with SNAP-Surface549). To observe the effect of DOR-Halo and KOR-Halo co-expression, the cells stably expressing Gqi5 and SNAPf-MOR were transiently transfected with the cDNAs encoding these proteins. The cells exhibiting spot densities of DOR-Halo or KOR-Halo at 0.75 ± 0.25 spots/µm^2^ (after labeling with Halo-SaraFluor650T) were selected for observations. Therefore, the Ca^2+^ mobilization assay was performed at about the same low expression levels of ORs as those employed for single fluorescent-molecule imaging experiments.

### Quantifying the internalization of SNAPf-ORs

See the companion paper under the same title.^1^

### Systemic morphine administration to mice and examination of its antinociceptive effect

Morphine hydrochloride (Takeda Pharmaceutical) was dissolved in saline and subcutaneously administered to mice once each day (10 mg/kg) for 11 days. Antinociception was measured by the tail-flick test on days 1, 5, 8, and 11 from 0 to 120 min after morphine administration (details will be described later). Body weight was measured before each morphine injection. The control group received saline without morphine. All animal experiments were randomized and performed by a blinded researcher, who was then unblinded before statistical analysis.

### Intracerebroventricular administration of the Dpep(20-42)DM peptide in mice

The Dpep(20-42)DM peptide was dissolved in freshly prepared aCSF (127 mM NaCl, 1.5 mM KCl, 1.24 mM KH_2_PO4, 1.4 mM MgSO_4_, 26 mM NaHCO_3_, 2.4 mM CaCl_2_, 10 mM D-glucose). For continuous intracerebroventricular administration, an osmotic minipump (Alzet model 2002) filled with the peptide solution was implanted according to the manufacturer’s protocol and as described previously.^80^ In order to stabilize the pump flow rate, the pumps were primed by placing them in sterile saline at room temperature overnight. The osmotic minipumps were filled with 200 µL of aCSF with or without the peptide solution (0.83 mg/mL in aCSF). The minipump was connected to a 2.5-mm-long cannula (Alzet Brain Infusion Kit 3) via a 1.5-cm-long polyvinylchloride tube. Before surgery, the mice were anesthetized with a combination anesthetic (0.75 mg/kg of medetomidine, 4.0 mg/kg of midazolam, and 5.0 mg/kg of butorphanol). The scalp was shaved, Povidone-Iodine (Isodine, Mundipharma K.K.) was applied, and an incision was made along the midline of the scalp; hemostats were then used to make a pocket under the skin between the shoulder blades. The skull was scraped clean of periosteum so that the cannula can properly adhere to the skull. A microdrill (Tamiya, Inc.) was used to create a hole approximately 1.1 mm lateral and 0.5 mm caudal to the bregma. The minipump was placed in the pocket under the skin between the shoulder blades, the cannula was inserted through the drilled hole into the lateral ventricle, and the cannula pedestal was affixed to the skull with cyanoacrylate glue and resin cement. The incision was closed with soft silk sutures and antibacterial ointment was applied to the wound. The animals were allowed to rest on a disposable heating pad until they were recovered by using 0.75 mg/kg of atipamezole and then returned to their home cages in the animal facility for 1-2 days until the measurement. Cannula placement into the lateral ventricle was verified with trypan blue (4%, 10 μl).

### Morphine-induced antinociception assay

Morphine-induced antinociception was evaluated by using the tail-flick test.^22^ Using a tail-flick apparatus (IITC Life Science), the intensity of the heat source was set at 10, which resulted in a basal tail-flick latency of 2-3 s for most of the animals. The tail-flick latency was recorded before (0 min, baseline latency) and every 30 min after morphine injection (30, 60, 90, and 120 min). The results are presented as a percentage of the maximal possible effect, %MPE, which was calculated for each mouse at each time point according to the following formula: %MPE = [(latency after drug − baseline latency)/ (10 − baseline latency)] × 100. Cutoff latency was selected at 10 s to minimize tissue damage. The area under the %MPE vs. time curve (AUC) for each treatment condition was also used to evaluate antinociception (Fig. 8h). Normal distribution of the data was verified before performing the parametric statistical analysis. Wherever appropriate, data were analyzed using two-way ANOVA, followed by Tukey’s post hoc tests or a multiple unpaired t-test with significance set at P < 0.05. All calculations were performed using the GraphPad Prism 9 software (GraphPad Software).

### Software and statistical analysis

The same as described in the companion paper.^1^

## Data availability

All data generated or analyzed for this study are available within the paper and its associated Supplementary Information. Any additional information required to reanalyze the data reported in this paper is available from the lead contact upon reasonable request. Superimpositions of image sequences obtained in two colors and tracking single-molecules for in-vitro observation data were performed using C + +-based computer programs, produced in-house and based on the well-established approaches.^81^ Further information regarding the experimental design may be found in the Nature Research Reporting Summary.

## Code availability

Source codes for this have been integrated into a large, complex software package. While the software package cannot be extracted in a useful way, the entire software is available from the lead contact upon reasonable request, who will provide guidance on how to use it, as its manual and comments are written in Japanese.

## Acknowledgements

We thank Profs. H. Takeshima of Kyoto University, R. Schülz of the University of Münich, and R. Y. Tsien of the University of California San Diego, for their kind gifts of cDNAs encoding rat KOR and DOR, that encoding rat MOR, and that encoding mCherry, respectively. We also thank Ms. Irina Meshcheryakova for technical help in preparing the cDNAs. This work was supported in part by Japan Society for the Promotion of Science (JSPS) Grants-in-Aid for Scientific Research (Kiban A to A.K. [21H04772], Kiban S to A.K. [16H06386], Kiban B to T.K.F. [16H04775, 20H02585], Kiban C to R.S.K. [17K07333], Wakate B to R.S.K. [26870292], Wakate to T.A.T. [21K15058], and Challenging Exploratory Research to T.K.F. [18K19001] and A.K. [22K19334]), a Grant-in-Aid from the Ministry of Education, Culture, Sports, Science and Technology of Japan (MEXT) for Transformative Research Areas (A) to T.K.F. (21H05252), and a JST grant ACT-X to T.A.T. (JPMJAX211B). WPI-iCeMS of Kyoto University is supported by the World Premiere Research Center Initiative (WPI) of MEXT.

## Contributions

P.Z., R.S.K., and A.K. conceived and formulated the project. A.K., T.A.T., T.K.F., and P.Z. developed the simultaneous dual-color single-molecule tracking station. P.Z. and A.K. designed biological experiments, and P.Z., with help from T.A.T., performed virtually all of the microscopic experiments. T.K.F., T.A.T., and A.K. developed and improved the in-house image analysis software and instrument control software, and discussed single-molecule imaging data. S.P. developed the theory for evaluating the dimer lifetimes from the single-molecule experimental data and assisted with its application to experimental results. T.A.T. and P.Z. analyzed the PCCF histograms and obtained dimer dissociation constants. W.F., with help from H.U., performed mouse experiments. P.Z., A.K., and T.A.T. wrote the manuscript, and all authors discussed the results and participated in the manuscript revisions.

