## Supplementary Information for "Single-molecule detection of transient dimerization of opioid receptors 2: Heterodimer blockage reduces morphine tolerance"

#### Captions to Supplementary Videos 1 and 2

#### Supplementary Figures 1-6

#### Supplementary Tables 1-5 (Supplementary Table 1 is an Excel table)

### Captions to Supplementary Videos 1

#### Supplementary Video 1

Simultaneous dual color imaging of single SNAPf-MOR (SNAP-Surface 549 label) and DOR-Halo7 (Halo7-SaraFluor650T label) molecules (both at densities of ≈0.8 spots/$\mu m^2$) in the PM at 37°C, recorded at video rate for 100 frames (3.3 s) and replayed at a 3x-slowed rate from real time. The movie on the right shows the results of the automatic detection of colocalized spots (yellow squares) in the movie on the left.

#### Supplementary Video 2

A typical colocalization event of two single SNAPf-MOR and DOR-Halo7 molecules in the PM (yellow circles) shown by expanded image frames, replayed at a 30x-slowed rate from real time.

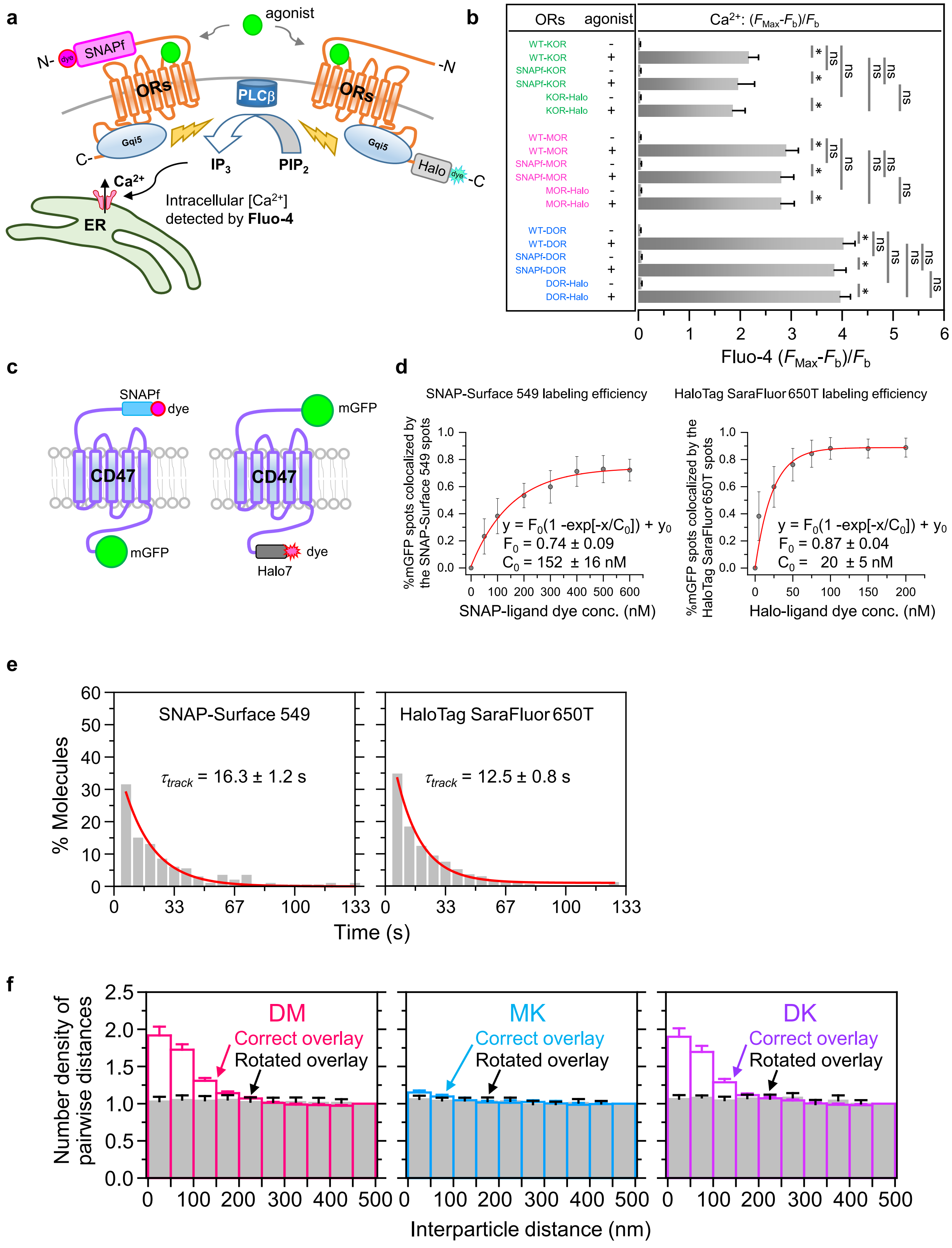


**Supplementary Fig. 1. Assessment of the functions of the SNAPf- and Halo-tagged ORs, labeling efficiencies of the tagged proteins, trackable duration lifetimes of the employed dye molecules, and PCCFs for the DM, MK, and DK heterodimers.**

(**a**) Schematic figure showing the experimental design for testing whether the ORs conjugated with the tag proteins SNAPf at the N-terminal ectodomain and Halo at the C-terminal endodomain retain signaling function comparable to non-tagged ORs, by observing $Ca^{2+}$ mobilization activated via Gqi5. For details, refer to Methods in this report, and STAR Methods and the caption to Supplementary Fig. 1a in the companion paper.

(**b**) $Ca^{2+}$ mobilization results showing that all three ORs fused with SNAPf- or Halo-tag proteins retain the functions of untagged ORs. $Ca^{2+}$ mobilization was parameterized by using $[F_{\mathrm{Max}}-F_{\mathrm{b}}]/F_{\mathrm{b}}$, where $F_{\mathrm{Max}}$ is the Fluo-4 peak signal intensity within 75 s after the addition of the stimulants and $F_{\mathrm{b}}$ is the baseline signal intensity.

(**c**) Evaluation method for the labeling efficiencies of the SNAPf-tag and Halo-tag proteins fused to the extracellular N-terminus or cytoplasmic C-terminus, respectively, of the monomeric TM protein CD47. SNAPf-CD47-mGFP and mGFP-CD47-Halo7 expressed in the PM were used. Since ORs form homodimers, their labeling efficiencies were difficult to evaluate, and therefore, the monomeric TM protein CD47 was employed for assessment purposes. Further details are provided in the companion paper.

(**d**) Determination of the extents of fluorescence labeling of SNAPf-OR and OR-Halo7 with the SNAP-Surface 549- and Halo-SaraFluor650T-conjugated ligands, respectively (mean ± SEM for n=20 cells). Each datapoint shows the percentage of fluorescent mGFP spots that were colocalized by the SNAP-dye spots (left) or Halo-dye spots (right) (mean ± SEM for n=20 cells), and these datapoints were plotted as a function of the dye concentration in the incubation medium (30-min reaction time). The plot could be fitted by a single exponential function plus a constant (values in the figure panels). Note that not every mGFP molecule was fluorescent, but since we examined whether the identified fluorescent mGFP molecule in the image was colocalized by a dye spot, the labeling efficiencies of the SNAPf- and Halo7-protein tags can be evaluated without the influence of the fluorescence efficiency of mGFP. The labeling efficiencies at 300 nM SNAP-Surface 549 and 100 nM Halo-SaraFluor650T, the conditions used throughout this study, were determined using the obtained exponential functions, providing efficiencies of 60% and 88% for SNAP-Surface 549 and Halo-SaraFluor650T, respectively.

(**e**) Distributions of the trackable duration times (trajectory lengths) of SNAP-Surface 549 and Halo-SaraFluor650T bound to SNAPf-MOR and MOR-Halo7, respectively, expressed at densities of ≈0.1 spots/$\mu m^2$ in CHO-K1 cells observed at 30 Hz. The decay time constants obtained by single exponential fitting provided the trackable duration lifetimes of the two fluorescent probes of 16.3 ± 1.2 for SNAP-Surface 549 and Halo-SaraFluor650T for 12.5 ± 0.8 s, under our experimental conditions. The photobleaching lifetimes for these fluorescent dye molecules adsorbed on the cover glass of the glass-base dish were 36.6 ± 1.0 s (n = 567) for SNAP-Surface 549 and 20.1 ± 0.10 s (n = 598) for SaraFluor650T.

(**f**) PCCFs for the DM, MK, and DK hetero-pairs of three ORs. Grey bars indicate the results of the 180°-rotated overlays of the simultaneously observed magenta and green images (control for incidental overlaps of fluorescent spots with different colors).

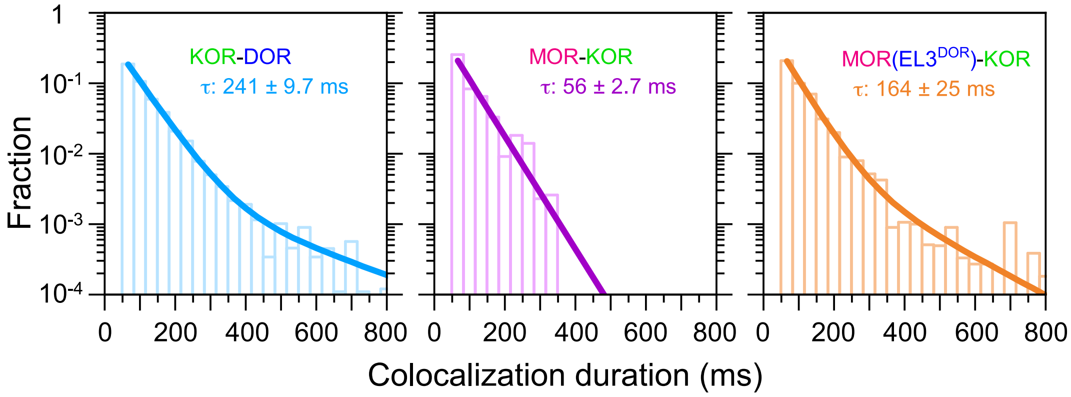


**Supplementary Fig. 2. Histograms showing the *duration* distributions of three different pairs of ORs, M($EL3^{DOR}$)-K, DK (positive control), and MK (negative control), indicating that $EL3^{DOR}$ is involved in DK heterodimerization.**

Histograms for the control pairs of DK and MK are the same as those shown in Fig. 1d.

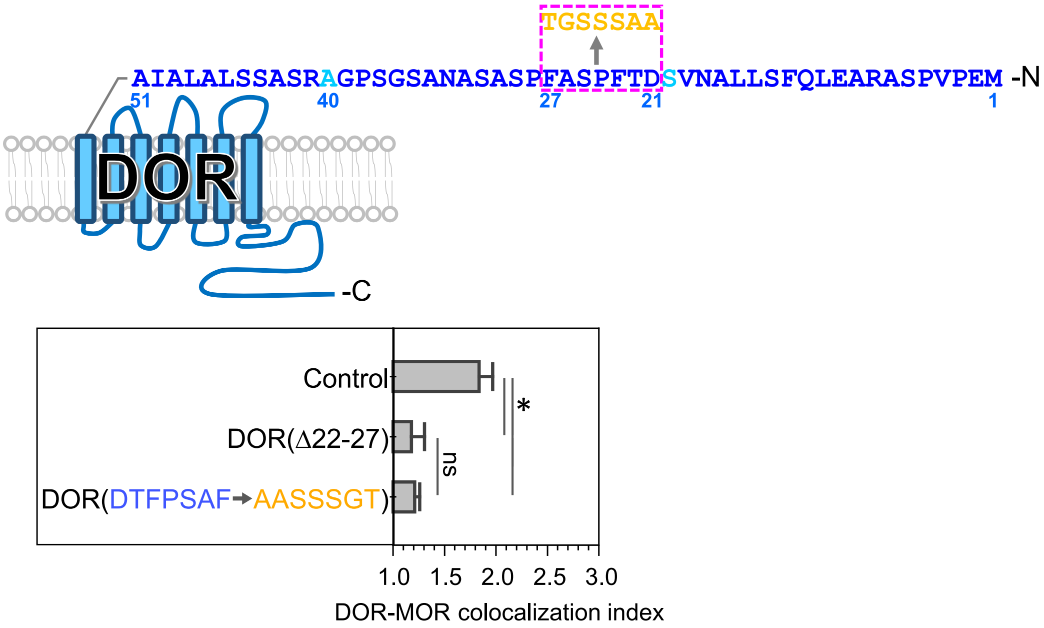


**Supplementary Fig. 3. Result showing that a DOR mutant with randomly substituted amino acids 21-27 (AASSSGT, an arbitrarily produced random sequence, replacing the original sequence of DTFPSAF; top illustrative figure) did not form detectable heterodimers with MOR (bottom bar graph). This result reinforces the data shown in Fig. 3a, which indicate that the deletion mutant DOR(Δ22-27) does not form heterodimers with MOR (bottom bar graph).**

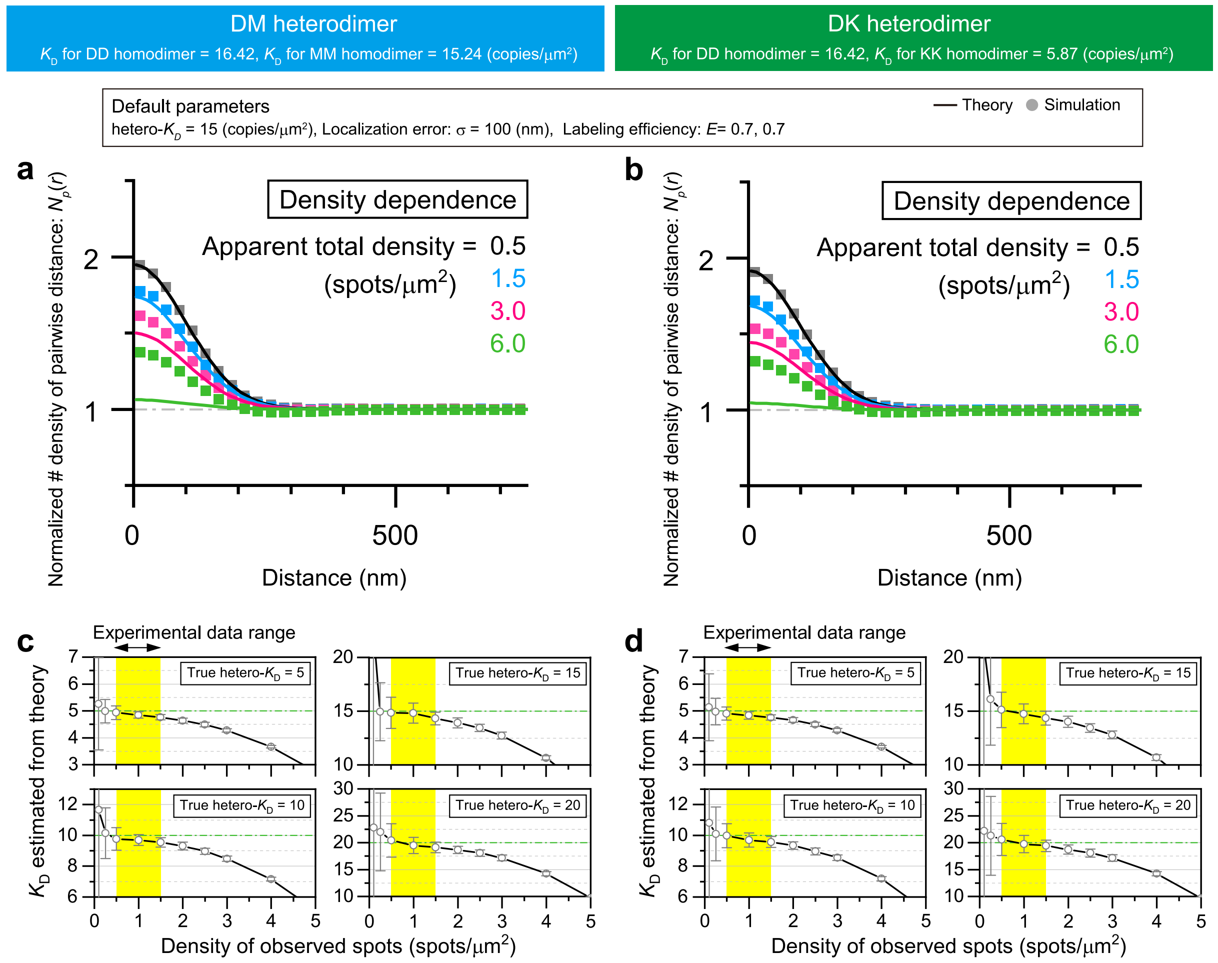


**Supplementary Fig. 4. Monte-Carlo simulation examination of the influences of the number density of fluorescent spots in the image (expression levels of molecules) on PCCF (a, b) and $K_D$ (c, d), obtained using the theory developed in Supplementary Note 2 in the companion paper.**

(**a**) and (**c**) are for DM heterodimers and (**b**) and (**d**) are for DK heterodimers.

For the details of (**a**) and (**b**), refer to Supplementary Fig. 3b, e and their captions, respectively, in the companion paper. The localization error found for DM and DK heterodimers (100 nm) is better than that found for homodimers (140 nm) described in the companion paper. This is probably due to the better signal-to-noise ratio we obtained for the Halo-SaraFluor650T used to observe heterodimers compared with that of the SNAP-CF660R used to observe homodimers.

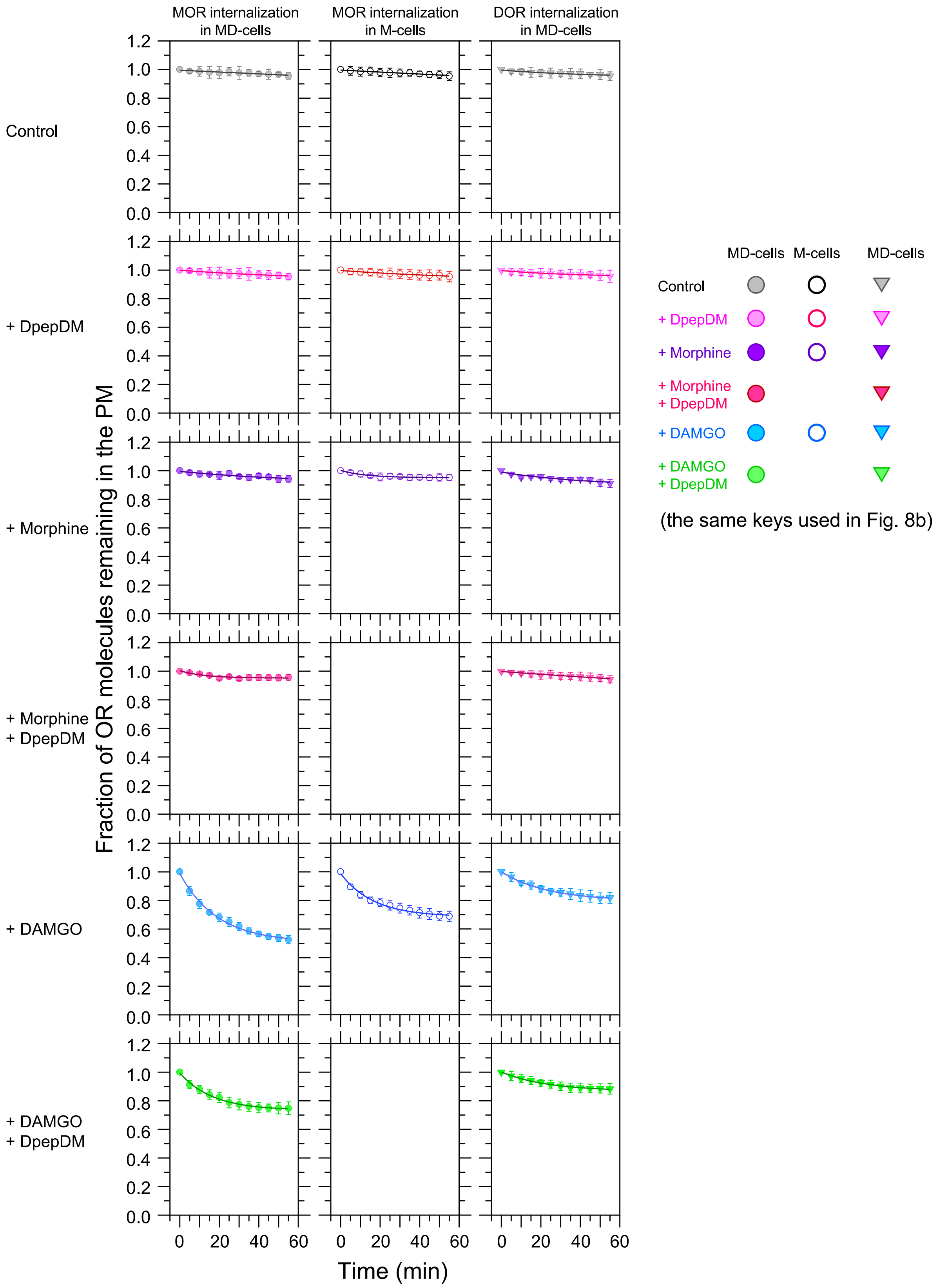


108

109 **Supplementary Fig. 5. Time course data for MOR and DOR internalization**
110 **obtained under different conditions are separately shown for enhanced**
111 **visibility of the datapoints and the best-fit curves.**

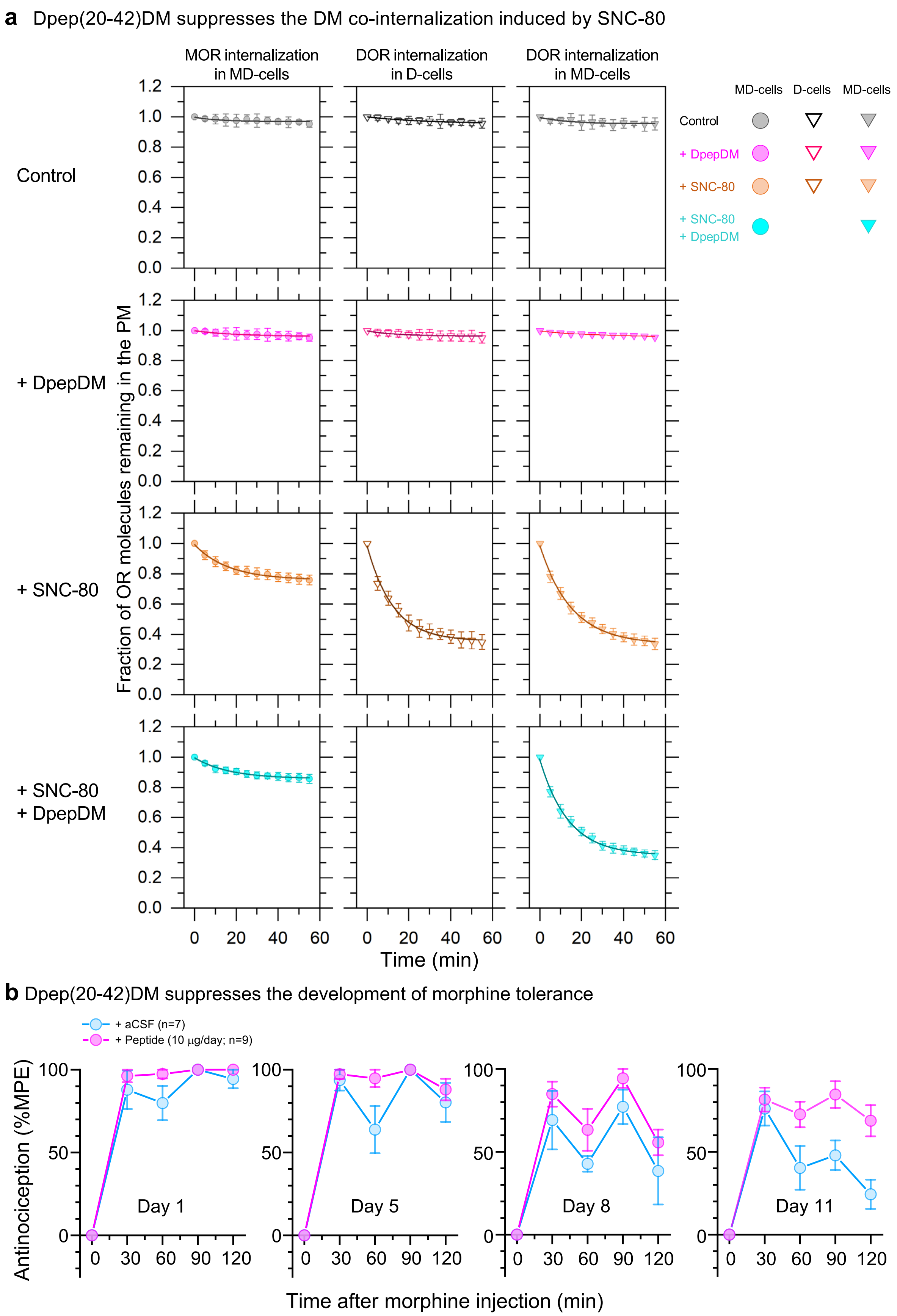


**Supplementary Fig. 6. Dpep(20-42)DM reduces D-agonist-induced MOR internalization in MD cells and morphine tolerance development in mice.**

(**a**) Dpep(20-42)DM reduces SNC-80 (D-agonist)-induced MOR internalization, but not DOR internalization in MD cells. Time courses of MOR and DOR internalization in CHO-K1 cells, before and after the addition of 0.5 μM SNC-80, a DOR agonist (top and third rows, respectively), and in the additional presence of 1 μM Dpep(20-42)DM (second and bottom rows). MOR internalization was examined only in MD-cells (left column), whereas DOR internalization was observed in D cells and MD-cells (middle and right columns, respectively). For the quantitative analysis method, see the caption to Fig. 8b. The summary of the fractions of internalized ORs, the residency lifetimes in the PM for the internalized fractions of molecules, and the statistical test results are shown in Supplementary Table 5.

(**b**) The antinociceptive effects of morphine (10 mg/kg/day) alone and in combination with Dpep(20-42)DM (10 μg/day) in an acute pain model using mice were examined by employing the tail flick test. The % maximal possible effect (%MPE) was obtained on days 1, 5, 8, and 11. All experiments were randomized and performed by a blinded researcher, who was then unblinded before statistical analysis.

**Supplementary Table 2. Summary of $K_D$, $k_{off}$, and $k_{on}$ for OR homodimers in the presence and absence of M-agonists, as well as the expected percentages of OR protomers existing as OR monomers and homo-dimers at various expression levels in the absence of other ORs.**

| | | | | Expected % protomers existing as dimers at different expression levels (copies/μm$^2$) | | | | |
|---|---|---|---|---|---|---|---|---|
| | $K_D$ (copies/μm$^2$) | $k_{off}$ (/s) | $k_{on}$ (μm$^2$/copies/s) | 0.3 Homo | 1 Homo | 3 Homo | 10 Homo | 30 Homo |
| KK homodimer | 5.87 ± 0.56 | 6.71 ± 1.17 | 1.14 ± 0.23 | 8.6 | 21.2 | 38.6 | 58.6 | 73.2 |
| DD homodimer | 16.42 ± 0.53 | 8.00 ± 0.96 | 0.49 ± 0.06 | 3.4 | 9.9 | 22.1 | 41.6 | 59.6 |
| + Morphine | 18.08 ± 1.44 | 8.13 ± 0.20 | 0.45 ± 0.04 | 3.1 | 9.1 | 20.8 | 39.9 | 58.1 |
| MM homodimer | 15.24 ± 1.54 | 8.47 ± 0.86 | 0.56 ± 0.08 | 3.7 | 10.5 | 23.2 | 42.9 | 60.7 |
| + Morphine | 19.69 ± 1.92 | 8.48 ± 0.22 | 0.43 ± 0.04 | 2.9 | 8.5 | 19.7 | 38.5 | 56.8 |
| + DAMGO | 8.34 ± 0.58 | 5.35 ± 1.69 | 0.64 ± 0.21 | 6.8 | 17.6 | 34.0 | 54.3 | 70.0 |

**Supplementary Table 3. Summary of $K_D$, $k_{off}$, and $k_{on}$ for DM and DK heterodimers in the presence and absence of M-agonists and Dpep(20-42)DM, as well as the percentages of OR protomers existing as OR homo- and hetero-dimers at various expression levels.**

The mean values of $K_D$, $k_{off}$, and $k_{on}$ are the same as those shown in Fig. 6e, but their SEMs are included in this table. The values before the M-agonist addition and after the DAMGO addition are the same as those shown in Fig. 4i of the companion paper, but the values after morphine addition are new. The expected percentages of OR protomers existing as homo- and hetero-dimers were calculated from the two homodimer $K_D$s and the heterodimer $K_D$. In this table, the expected percentages are only shown for the cases where the expression levels of DOR and MOR or DOR and KOR are the same (for example, 3 + 3 indicates the case where both DOR and MOR are expressed at 3 copies/μm$^2$). The results when the expression levels of DOR and MOR or DOR and KOR are different are shown in Fig. 7 (only in the absence of DpepDM). The percentages of OR protomers existing as homo- and hetero-dimers are correct only in the absence of other ORs and other unknown molecules competing for the same binding sites of ORs (which might or might not exist in the CHO-K1 cells employed here).

| | | $K_D$ (copies/μm$^2$) | $k_{off}$ (/s) | $k_{on}$ (μm$^2$/copies/s) | Expected % protomers existing as dimers at different expression levels (copies/μm$^2$): 0.3 + 0.3 | | | 1 + 1 | | | 3 + 3 | | | 10 + 10 | | | 30 + 30 | | |
|---|---|---|---|---|---|---|---|---|---|---|---|---|---|---|---|---|---|---|---|
| | | | | | MM | DD | DM | MM | DD | DM | MM | DD | DM | MM | DD | DM | MM | DD | DM |
| DOR + MOR | | | | | | | | | | | | | | | | | | | |
| No addition | | 14.00 ± 1.59 | 3.85 ± 0.18 | 0.28 ± 0.03 | 1.8 | 1.6 | 1.9 | 4.8 | 4.5 | 5.2 | 9.7 | 9.2 | 10.6 | 16.2 | 15.7 | 18.0 | 21.5 | 21.0 | 24.0 |
| | + DpepDM | 36.53 ± 7.12 | 8.07 ± 0.72 | 0.22 ± 0.04 | 1.8 | 1.7 | 0.8 | 5.1 | 4.8 | 2.1 | 10.8 | 10.3 | 4.5 | 19.0 | 18.4 | 8.1 | 26.2 | 25.7 | 11.2 |
| + Morphine | | 15.36 ± 3.00 | 5.35 ± 0.57 | 0.35 ± 0.07 | 1.5 | 1.4 | 1.8 | 4.2 | 3.9 | 4.9 | 8.7 | 8.2 | 10.3 | 15.0 | 14.4 | 18.0 | 20.3 | 19.7 | 24.6 |
| | + DpepDM | 23.65 ± 10.08 | 8.13 ± 0.60 | 0.34 ± 0.14 | 1.5 | 1.4 | 1.2 | 4.3 | 4.0 | 3.3 | 9.2 | 8.7 | 7.1 | 16.4 | 15.7 | 12.8 | 22.6 | 22.1 | 17.8 |
| + DAMGO | | 6.94 ± 0.72 | 2.71 ± 0.08 | 0.39 ± 0.04 | 2.9 | 1.6 | 3.6 | 7.1 | 4.2 | 9.1 | 12.3 | 8.2 | 16.9 | 17.7 | 13.5 | 26.1 | 21.3 | 17.9 | 33.0 |
| | + DpepDM | 16.20 ± 2.49 | 5.65 ± 0.48 | 0.35 ± 0.05 | 3.1 | 1.7 | 1.6 | 7.7 | 4.6 | 4.3 | 14.3 | 9.6 | 8.4 | 21.8 | 16.8 | 13.8 | 27.2 | 23.1 | 18.1 |
| | | | | | KK | DD | DK | KK | DD | DK | KK | DD | KD | KK | DD | DK | KK | DD | DK |
| DOR + KOR | | | | | | | | | | | | | | | | | | | |
| No addition | | 14.76 ± 1.73 | 4.15 ± 0.17 | 0.28 ± 0.03 | 4.1 | 1.6 | 1.7 | 9.8 | 4.6 | 4.4 | 17.0 | 9.6 | 8.5 | 24.4 | 16.9 | 13.5 | 29.3 | 23.4 | 17.4 |
| | + DpepEL3 | 29.30 ± 3.52 | 6.10 ± 0.93 | 0.21 ± 0.03 | 4.2 | 1.7 | 0.9 | 10.2 | 4.7 | 2.3 | 18.0 | 10.3 | 4.6 | 26.5 | 18.6 | 7.4 | 32.5 | 26.2 | 9.8 |

**Supplementary Table 4. Summary of the percentages of OR protomers at various expression levels existing as OR monomers and homo- and hetero-dimers in the presence and absence of M-agonists and Dpep(20-42)DM.**

The expected percentages of DOR or MOR protomers existing as monomers and homo- and hetero-dimers, when DOR and MOR are expressed at approximately equal number densities, were calculated from the two homodimer $K_D$s and the heterodimer $K_D$ (for example, 3 + 3 indicates the case where both DOR and MOR are expressed at 3 copies/μm$^2$). The results when the expression levels of DOR and MOR are different are shown in Fig. 7 (only in the absence of DpepDM). The percentages are only correct in the absence of other ORs and other unknown molecules competing for the same binding sites of ORs (which might or might not exist in the CHO-K1 cells employed here).

Expected % MOR protomers existing as monomers and dimers at different expression levels (copies/μm$^2$)

| | | 0.3 + 0.3 | | | 1 + 1 | | | 3 + 3 | | | 10 + 10 | | | 30 + 30 | | |
|---|---|---|---|---|---|---|---|---|---|---|---|---|---|---|---|---|
| | | M | MM | DM | M | MM | DM | M | MM | DM | M | MM | DM | M | MM | DM |
| DOR + MOR | | | | | | | | | | | | | | | | |
| No addition | | 94.6 | 3.5 | 1.9 | 85.2 | 9.5 | 5.2 | 70.0 | 19.3 | 10.6 | 49.7 | 32.4 | 18.0 | 33.0 | 43.0 | 24.0 |
| | + DpepDM | 95.6 | 3.6 | 0.8 | 87.8 | 10.1 | 2.1 | 73.9 | 21.5 | 4.5 | 53.8 | 38.0 | 8.1 | 36.5 | 52.3 | 11.2 |
| + Morphine | | 95.2 | 3.0 | 1.8 | 86.7 | 8.3 | 4.9 | 72.3 | 17.3 | 10.3 | 52.0 | 29.9 | 18.0 | 34.9 | 40.5 | 24.6 |
| | + DpepDM | 95.8 | 3.0 | 1.2 | 88.1 | 8.6 | 3.3 | 74.5 | 18.4 | 7.1 | 54.4 | 32.8 | 12.8 | 36.9 | 45.3 | 17.8 |
| + DAMGO | | 90.5 | 5.9 | 3.6 | 76.7 | 14.1 | 9.1 | 58.5 | 24.6 | 16.9 | 38.5 | 35.5 | 26.1 | 24.4 | 42.7 | 33.0 |
| | + DpepDM | 92.3 | 6.1 | 1.6 | 80.3 | 15.5 | 4.3 | 63.0 | 28.6 | 8.4 | 42.6 | 43.6 | 13.8 | 27.5 | 54.4 | 18.1 |

Expected % DOR protomers existing as monomers and dimers at different expression levels (copies/μm$^2$)

| | | 0.3 + 0.3 | | | 1 + 1 | | | 3 + 3 | | | 10 + 10 | | | 30 + 30 | | |
|---|---|---|---|---|---|---|---|---|---|---|---|---|---|---|---|---|
| | | D | DD | DM | D | DD | DM | D | DD | DM | D | DD | DM | D | DD | DM |
| DOR + MOR | | | | | | | | | | | | | | | | |
| No addition | | 94.8 | 3.3 | 1.9 | 85.8 | 9.0 | 5.2 | 71.0 | 18.4 | 10.6 | 50.7 | 31.3 | 18.0 | 33.9 | 42.1 | 24.0 |
| | + DpepDM | 95.9 | 3.4 | 0.8 | 88.4 | 9.5 | 2.1 | 74.9 | 20.5 | 4.5 | 55.0 | 36.9 | 8.1 | 37.5 | 51.3 | 11.2 |
| + Morphine | | 95.4 | 2.8 | 1.8 | 87.3 | 7.7 | 4.9 | 73.3 | 16.4 | 10.3 | 53.2 | 28.8 | 18.0 | 36.0 | 39.5 | 24.6 |
| | + DpepDM | 96.0 | 2.8 | 1.2 | 88.7 | 8.0 | 3.3 | 75.5 | 17.4 | 7.1 | 55.7 | 31.5 | 12.8 | 38.1 | 44.1 | 17.8 |
| + DAMGO | | 93.2 | 3.2 | 3.6 | 82.6 | 8.3 | 9.1 | 66.8 | 16.3 | 16.9 | 47.0 | 26.9 | 26.1 | 31.3 | 35.8 | 33.0 |
| | + DpepDM | 95.1 | 3.3 | 1.6 | 86.6 | 9.1 | 4.3 | 72.4 | 19.2 | 8.4 | 52.5 | 33.6 | 13.8 | 35.6 | 46.3 | 18.1 |

**Supplementary Table 5. Summary of the fractions of internalized ORs, detectable by an observation duration of 55 min, and their residency lifetimes in the PM. The agonists (0.5 μM) were morphine and DAMGO for MOR and SNC-80 for DOR.**

| OR and Cells | Additions | MOR | | DOR | |
|---|---|---|---|---|---|
| | | Fraction of internalized molecules (%) | Residency time of internalized molecules (min) | Fraction of internalized molecules (%) | Residency time of internalized molecules (min) |
| MOR in M cells | Control | 3.5 ± 0.7 | 20.0 ± 11 | - | - |
| | + DpepDM | 4.2 ± 0.7 | 26.3 ± 11 | - | - |
| | + Morphine | 5.5 ± 0.3 | 21.8 ± 3.5 | - | - |
| | + DAMGO | 30.2 ± 1.1* | 16.6 ± 1.7* | - | - |
| DOR in D cells | Control | - | - | 4.4 ± 0.5 | 26.4 ± 8.0 |
| | + DpepDM | - | - | 4.0 ± 0.6 | 25.2 ± 9.6 |
| | + SNC-80 | - | - | 62.8 ± 1.6* | 12.4 ± 0.8* |
| DOR & MOR In MD cells | Control | 3.1 ± 0.7 | 21.0 ± 12 | 4.5 ± 0.8 | 20.8 ± 9.9 |
| | + DpepDM | 3.7 ± 0.6 | 20.0 ± 8.5 | 4.0 ± 0.5 | 27.4 ± 9.4 |
| | + Morphine | 5.7 ± 1.4 | 28.1 ± 17 | 7.3 ± 1.1 | 20.6 ± 9.3 |
| | + Morphine & DpepDM | 6.4 ± 0.3 | 26.0 ± 3.5 | 5.2 ± 1.1 | 28.5 ± 15 |
| | + DAMGO | 48.1 ± 1.2* | 18.9 ± 1.3 | 20.3 ± 0.5* | 23.0 ± 1.5 |
| | + DAMGO & DpepDM | 26.0 ± 0.7*† | 16.5 ± 1.3* | 13.3 ± 0.5*† | 23.2 ± 2.2 |
| | + SNC-80 | 23.3 ± 0.6* | 16.1 ± 1.2* | 65.2 ± 1.2* | 15.3 ± 0.8* |
| | + SNC-80 & DpepDM | 13.9 ± 0.5*‡ | 16.4 ± 1.6* | 66.8 ± 1.2* | 15.4 ± 0.6* |

*: Significantly different from respective control specimens. Similar comparisons with respective control specimens were made for all the cases, but no significant differences were detected other than those indicated with * (no indication is given within the table).

† and ‡: Significantly different from "+ DAMGO" and "+ SNC-80", respectively. The effects of DpepDM were undetectable except for the cases shown by † and ‡.
